# Structural Insights into Negative Cooperativity between Gemin2 and RNA in Sm-class snRNP Assembly

**DOI:** 10.1101/857334

**Authors:** Rundong Zhang

**Affiliations:** Department of Ophthalmology, State Key Laboratory of Biotherapy, West China Hospital, Sichuan University and Collaborative Innovation Center, Chengdu 610041, P. R. China

## Abstract

Sm-class ribonucleoprotein particles (RNPs) are ring-shaped structures (Sm cores) formed by Sm hetero-heptamer around a segment of RNA, containing a nonameric oligoribonucleotide, PuAUUUNUGPu, followed by a stem-loop, and are basic structural modules critical for stability and functions of spliceosomal, telomerase and U7 RNPs. In the chaperones-assisted Sm core assembly, Gemin2 of the SMN complex, not only binds SmD1/D2/F/E/G (5Sm), but also serves as a checkpoint via a negative cooperativity mechanism uncovered in our recent study: Gemin2 constricts the horseshoe-shaped 5Sm in a narrow conformation from outside, preventing non-cognate RNA and SmD3/B from joining; only cognate RNA can bind inside 5Sm and widen 5Sm, dissociating Gemin2 from 5Sm and recruiting SmD3/B. However, the structural mechanics is unknown. Here I describe a coordinate-improved structure of 5Sm bound by Gemin2/SMN. Moreover, via new analysis, comparison of this structure with those of newly coordinate-improved Sm cores reveals the negative cooperativity mechanism between Gemin2 and RNA in binding 5Sm at atomic resolution level and provides structural insights into RNA selection and Gemin2’s release in Sm core assembly. Finally, implications in the evolution of the Sm-core assembly chaperoning machinery and the neurodegenerative disease spinal muscular atrophy caused by SMN deficiency are discussed.

## INTRODUCTION

Sm-class ribonucleoprotein particles (RNPs) are a major class of non-coding RNA-protein complexes in eukaryotes that form basic modules for more complicated RNA-protein complexes and play key roles in gene posttranscriptional expression, including pre-messenger RNA splicing [mediated by major spliceosomes U1, U2, U4, U5 and minor spliceosomes U11, U12 and U4atac small nuclear RNPs(snRNPs)] (1–3), suppression of premature cleavage (U1 snRNPs) (4) and histone RNA 3’-end processing (U7 snRNPs) (5,6), as well as in maintenance of chromosome ends (telomerase RNA subunit) (7,8). They are also found in viral RNAs and play important roles in viral functions (9,10). The first and also most studied Sm-class RNPs are the cores of the above-mentioned spliceosomal snRNPs, termed as snRNP cores or Sm cores. In the Sm core, seven Sm proteins—D1, D2, F, E, G, D3, and B—in the order, form a ring-like structure around a special segment of each snRNA, termed as snRNP code, consisting of a nonameric oligoribonucleotide (Sm site), usually PuAUUUNUGPu, plus an adjacent 3’-stem-loop (SL), which usually contains at least 6-7 base pairs (11). U7 snRNP cores are special Sm-class RNPs, because SmD1/D2 is replaced by heterodimeric Sm-like protein Lsm10/11 and a special Sm site is used, which is AAUUUGU<u>CUAG</u> (the difference from the above-mentioned Sm site is underlined)(6,12). However, U7 snRNP cores follow the same maturation pathway as the spliceosomal Sm-class snRNP cores do (5,6,13). Therefore I will discuss their assembly together with the spliceosomal Sm cores as Sm-class RNP assembly or Sm core assembly.

The assembly of Sm cores is a key step in snRNP biogenesis. It takes a stepwise fashion as has been demonstrated by *in vitro* study from three heteromeric Sm subcomplexes, SmD1/D2, SmF/E/G, and SmD3/B (14,15). At first, SmD1/D2 and SmF/E/G associate to form a metastable intermediate complex, which then binds RNA to form a stable Sm subcore. At last, SmD3/B joins to form a highly stable Sm core. The reaction *in vitro* is a spontaneous process and Sm cores can form on any RNA containing just the nonameric Sm site (14,15). However, in eukaryotic cells (especially more complexed metazoans), two complexes, the PMRT5 (protein arginine methyltransferase 5) and SMN (survival motor neuron) complexes, are sequentially involved in Sm core assembly and they help enhance the assembly specificity of Sm proteins, allowing them to assemble only on the cognate RNAs, mostly snRNAs, which contain not only the Sm site, but a 3’-adjacent stem-loop as well (11,16–18).

The PRMT5 complex is comprised of PRMT5, WD45 and pICln. PRMT5 and WD45 form a hetero-octamer and methylate the C-terminal arginine residues of SmD3, SmB and SmD1, which is thought to enhance the interactions between Sm proteins and SMN (19,20). pICln binds SmD1/D2 and SmF/E/G into a ring-shaped 6S complex, stabilizing these two Sm subcomplexes in the finally assembled order and simultaneously preventing the entry of any RNA to the RNA-binding pocket (17). In addition, pICln also binds SmD3/B (17).

The SMN complex consists of 9 proteins—SMN, Gemin2-8 and unrip—in higher eukaryotes and functions in the later phase. It accepts SmD1/D2/F/E/G (5Sm) and SmD3/B and releases pICln (17), and also ensures that Sm proteins assemble only on cognate snRNAs (11,18). Gemin2 is the most conserved component in the SMN complex as only its orthologues are found in all eukaryotes, including unicellular eukaryotes, like *S. cerevisiae* (21,22). Gemin2 is the acceptor of 5Sm (23,24), which binds the horseshoe-shaped 5Sm from the peripheral side, with its N- and C-terminal domains (NTD and CTD) binding F/E and D1/D2 respectively, and leaves the RNA-binding channel of 5Sm open for RNA to enter (23). SMN is the second conserved component (21,22) and seems to serve as a scaffold protein. It tightly binds Gemin2 by its N-terminal Gemin2-binding domain (Ge2BD, residues 26-51) (23,25) and also interacts with Gemin8 by its C-terminal self-oligomized YG box (26). In vertebrates, either *SMN* or *Gemin2* gene knockout causes early embryonic death, indicating their essential roles in eukaryotic cells (27,28). Moreover, the deficiency of SMN causes neurodegenerative disease spinal muscular atrophy (SMA), emphasizing the medical relevance of the Sm-core assembly pathway (29–31). Gemin8 further binds Gemin6/7 and Unrip, but their roles are poorly understood (26,32,33). Gemin3, a putative RNA helicase (34), associates with Gemin4 (35), and both are required for Sm core assembly in higher eukaryotes, but their specific functions are unknown. Gemin5 is the component to initially bind precursor (pre)-snRNAs and deliver them to the rest of the SMN complex for assembly into the Sm core (36). Although it had long been considered as the protein conferring the RNA assembly specificity by direct recognition of the snRNP code (37–42), our recent study indicates that it is Gemin2 that plays the role of enhancing RNA assembly specificity by an unusual yet elegant way—via negative cooperativity with RNA in binding to 5Sm (43).

Gemin2, independent of its N-terminal tail (residues 1-39), constricts the horseshoe-shaped 5Sm in a narrow conformation by binding its outside (43). The nonameric Sm site RNA, AAUUUUUGA, which can form the Sm subcore if incubated with SmD1/D2 and SmF/E/G alone, cannot bind 5Sm again when it is bound by Gemin2, even if additional RNA sequence is at its 5’-side (43). Only the RNAs with a 3’-SL immediately following the Sm site can preferentially get into the RNA-binding pocket inside the horseshoe-shaped 5Sm, but only to a small extent (43). The binding of RNAs obviously widens the opening between SmD1 and SmG, which makes space for SmD3/B to bind to form the final Sm core. Moreover, the widening of SmD1-SmG also causes Gemin2 to detach from 5Sm and the joining of SmD3/B facilitates it (43) (Figure 1A). As there is no spatial clash between Gemin2 and RNA’s binding to 5Sm, but each’s binding to 5Sm inhibits the other’s, Gemin2 and RNA are in the relationship of negative cooperativity in binding to 5Sm (43). This finding provides insights into these two basic questions: how the SMN complex enhances RNA assembly specificity and how it releases (43). However, in terms of the structural point of view, these questions are yet not answered at atomic resolution level: How does Gemin2’s constriction of 5Sm in a narrow conformation enhance RNA assembly specificity? How does RNA binding cause the conformation of 5Sm to change? How does 5Sm change its conformation? How does the conformational change of 5Sm cause Gemin2 to release from the Sm subcore? And how does the joining of SmD3/B facilitate it?

**Figure 1.**
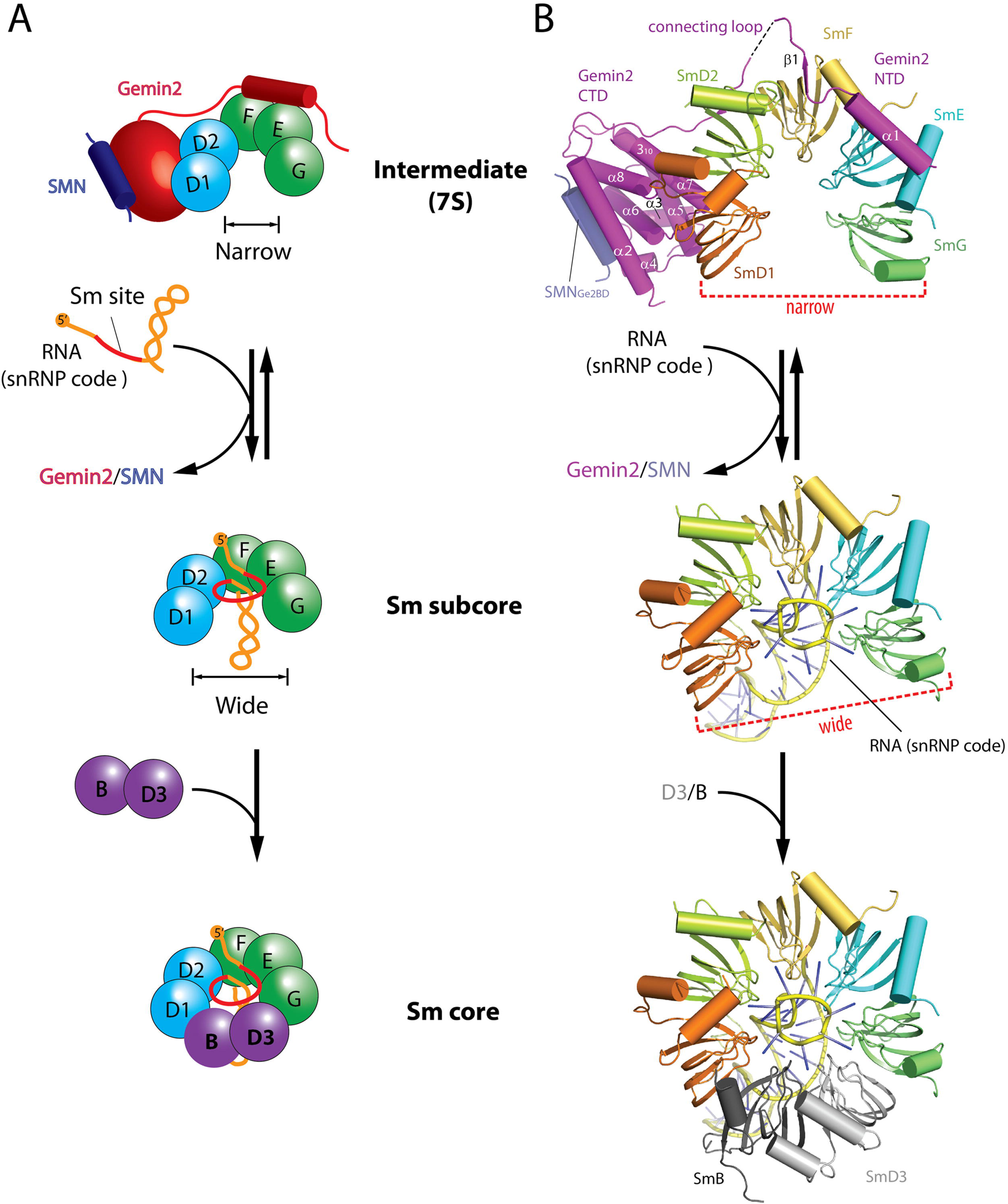
The transition during Sm core assembly from the 7S intermediate to the Sm subcore and to the Sm core. The process is represented in cartoon (A) and structure (B). In the coordinate-improved structure of the 7S intermediate complex (PDB code 5XJL), for clarity, the N-tail of Gemin2 (residues 1-47, only 22-31 is visible in the crystal structure) is not shown. The Sm subcore, which has no crystal structure determined, is represented by the portion of the U4 Sm core (PDB code 4WZJ)(45) without SmD3/B. The mature Sm core is represented by U4 snRNP (PDB code 4WZJ). For clarity, the preceding 5’-segment before the Sm site in U4 snRNA is not shown.

To address these questions, at first, I describe a re-refined and coordinate-improved structure of the key intermediate of the SMN complex we had previously determined (23), the 7S complex, which contains Gemin2/SMN (residues 26-62)/SmD1/D2/F/E/G. Second, using a new method different from our previous (23), I made comparison and analysis of the new 7S structure and two recent structures of mature snRNP cores with improved resolution and coordinates (44,45). Based on them, I propose a model at atomic resolution level for the series of conformational changes from the 7S complex to the Sm subcore and to the Sm core, and discuss the structural basis for RNA selection and Gemin2’s release. Finally, with integration of the new understandings of Sm core assembly at the biochemical level (43) as well as the structural level (uncovered in this study), implications in the evolution of the chaperoning machinery for Sm core assembly and in SMA are discussed.

## MATERIALS AND METHODS

### Re-refinement of the crystal structure of the 7S complex

The previously determined crystal structure of the 7S complex (23) is re-refined with reference to two related complex structures accessible in recent years, the NMR structure of the complex Gemin2 (residues 95-280)/SMN (residues 26-51) (PDB code 2LEH) (25) and the 8S complex (PDB code 4V98), containing human SmD1/D2/F/E/G and *Drosophila melanogaster* pICln (residues 1-180), SMN (residues 1-122) and full-length Gemin2 (residues 1-245) (24) by cycles of manual rebuilding in Coot (46) and REFMAC refinement (47) in CCP4 suite (48). The final structure has improved quality as indicated by the reduction of R and R_free_ from 25.3% and 33.2% to 21.7% and 29.5% respectively (Supplementary Table S1). The final model (PDB code 5XJL) contains SmD1 (residues 1-81), SmD2 (residues 21-77 and 89-117), SmF (residues 3-76), SmE (residues 14-90), SmG (residues 8-52 and 55-72), Gemin2 (residues 22-31, 48-73, 79-124, 135-151, and 174-278), and SMN (residues 35-51). Ramachandran plot shows 95.8% of the dihedral angles in favored region, 2.6% in additional allowed region, and 1.6 (9 out of 576) in disallowed region (Supplementary Table S1). All the 9 outliers are located in the loop regions with relatively poor electron density. Only the regions supported by high quality electron density maps are presented and discussed in detail.

### Building of the Sm site RNA model in the 7S complex

The first 7 nucleotides of the Sm site in U4 snRNP (PDB code 4WZJ) (45) were individually saved together with their interacting Sm proteins. Each of the coordinates was then aligned with its corresponding Sm protein in the 7S complex. The 7 nucleotides were linked and followed by a relaxing of conformational constrains in Coot (46).

### Multiple sequence alignment

Multiple sequence alignments for Gemin2 and Sm/Lsm proteins in various species were performed by Clustal Omega (49) and the figures were prepared by ESPript (50).

### Structure alignment, analysis and preparation of images

PyMOL (The PyMOL Molecular Graphics System, Version 1.3, Schrödinger) was used for structure alignment and analysis as well as preparation of images.

## RESULTS AND DISCUSSION

### The refined structure of the SmD1/D2/F/E/G/Gemin2/SMN (residues 26-62) complex

We previously resolved the crystal structure of the key intermediate of the SMN complex, the SmD1/D2/F/E/G/Gemin2/SMN(residues 26-62) complex (the 7S complex for brevity), to a resolution at 2.5 Å at one direction (23). Although majority of the structure is accurate, due to its anisotropic feature at high resolution, some parts of the complex structure were not precisely determined, which hinders analysis and interpretation of the mechanics of conformational changes of the 7S complex, especially on Gemin2’s release caused by RNA-dependent 5Sm splay. In addition, in recent years, two related complex structures, the NMR structure of the complex Gemin2 (residues 95-280)/SMN (residues 26-51) and the 8S complex, containing human SmD1/D2/F/E/G and *Drosophila melanogaster* pICln (residues 1-180), SMN (residues 1-122) and full-length Gemin2 (residues 1-245), were determined (24,25), which makes it possible to improve the 7S coordinate with reference to them. In this study, I reprocessed the 7S complex data and gained a better quality of the 7S coordinate as indicated by the reduction of R and R_free_ (Supplementary Table S1). In the new structure (Figure 1B), there is little change in the 5 Sm proteins and the SMN segment, but a few subtle yet significant improvements in Gemin2, which are critical for interpretation of the structural mechanism of negative cooperativity between Gemin2 and RNA in binding to 5Sm. One significant place is in the N-terminal E/F binding domain of Gemin2, where the linker (residues 63-65) between α1 and β1 is improved (details are described below). The second place is Gemin2’s C-terminal tail (residues 270-280), where a 3_10_-helix is rebuilt (details are described below). Both these improvements are critical to the explanation of Gemin2-5Sm interactions and Gemin2’s release from 5Sm upon RNA binding. Another significant place is α3 (residues 139-151), where the register error is corrected, which contributes to the reduction of R values, although this correction plays little role in Gemin2-5Sm interactions.

As described before (23), the overall architecture of the 7S complex is that the 5 Sm proteins, D1, D2, F, E and G, form a horseshoe-shaped heteropentamer, with Gemin2 bound on the periphery of 5Sm by its two major domains NTD and CTD (Figure 1B). Connecting the NTD and CTD is a long loop, which is flexible as indicated by the significant loss of electron density. Although in the previous study the N-terminal tail of Gemin2 resides inside the RNA-binding pocket of 5Sm and was thought to play an inhibitory role in RNA binding (23), our recent characterization showed that the N-tail can automatically flip out of the RNA-binding pocket and seems to play a less significant role in RNA selection (43), and therefore it is not drawn in Figure 1B and will not be discussed below.

At the interface of Gemin2-SmE/F (Figure 2A-B), most of the interactions have been described in our previous study (23). However, new interactions of the linker (residues 63-65) between α1 and β1 of Gemin2 to SmF are found (Figure 2B). The last 2 residues in the 3-residue linker provide two carbonyl oxygens from the main chain to form a hydrogen bonding network with the side chains of Lys8 and Asn12 on the N-terminal helix of SmF. In this way, the tight hydrogen bonding interactions are continuous from the main chain of Pro64 to Ala69 of Gemin2, like a zipper, covering last two thirds of the linker and the entire following β1. Gemin2’s α1 has two types of interactions to SmE/F: the first one is on the N-terminal part of α1, where the hydrophobic interactions dominate between Pro49, Tyr52, Leu53 and Val56 from Gemin2, and Ile18, Phe22, Leu25 and Phe50 from SmE. These interactions are kind of "ridges-into-grooves" pattern. The second type is hydrogen bonding interactions from three residues, Tyr52, Gln57 and Glu59 from the middle part of Gemin2’s α1, to the main chain amide of Phe50 and the side chain of Glu52 from SmE, and to the side chain of Asn6 from SmF, respectively. Overall, Gemin2’s NTD is like two sticks, one being α1 (residues 49-62) and the other being the "zipper" (residues 64-69), connected by a short joint (residue 63). Roughly each "stick" interacts with one helix and one strand from each Sm protein.

**Figure 2.**
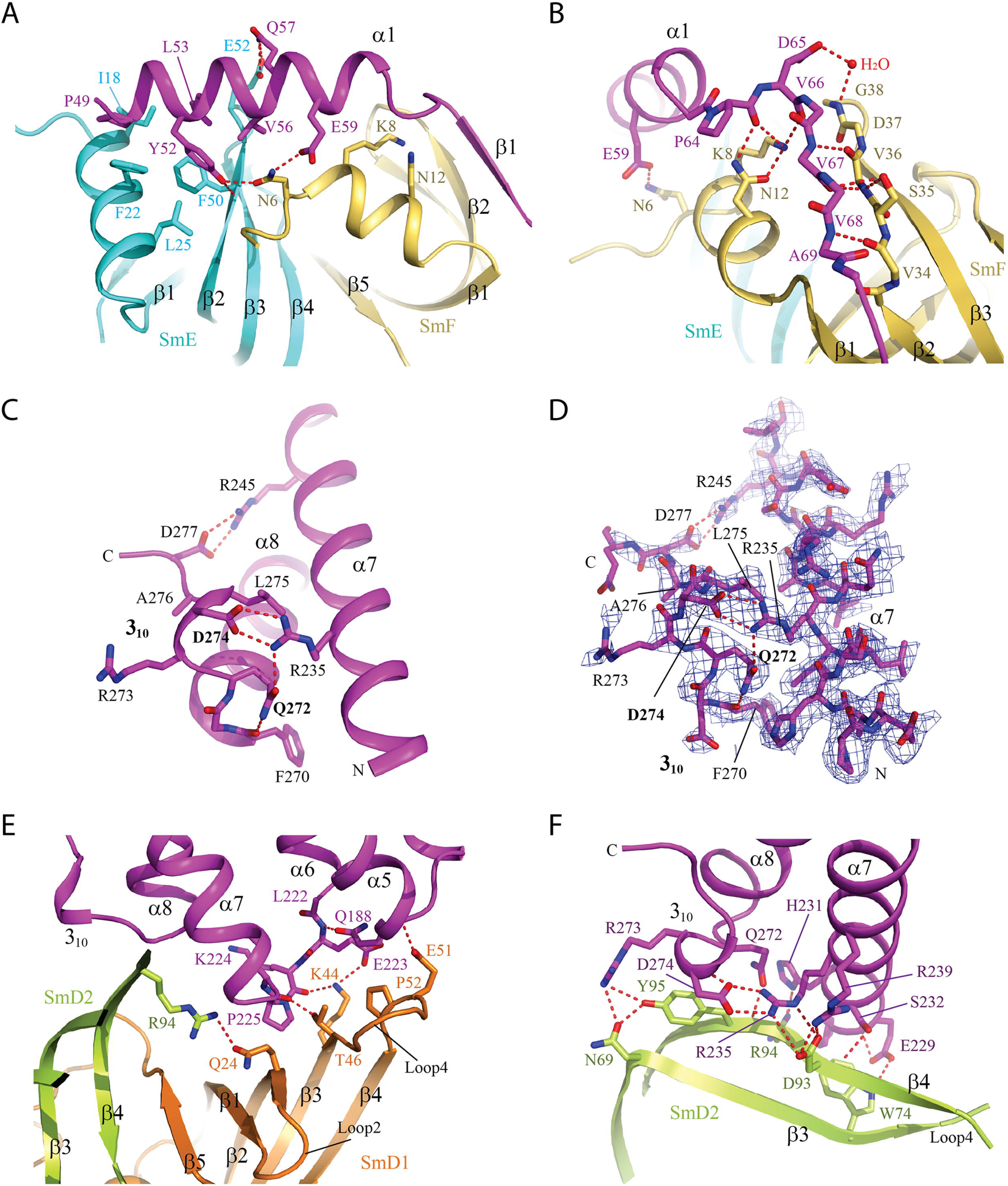
Detailed views of interactions between Gemin2 and Sm proteins. (A-B) Gemin2’s NTD interacts with SmF/E. (C-D) the C-terminal 3_10_ helix of Gemin2 and its interactions with helices α7-8. SigmaA-weighted 2Fo-Fc electron density map (blue mesh) is contoured at 1.1σ in (D). (E-F) Gemin2’s CTD interacts with SmD1/D2. Color scheme is the same as in Figure 1B. Hydrogen bonds and salt bridges are shown as red dashed lines. Oxygen and nitrogen are colored in red and blue respectively. Water is shown in red sphere. See main text for detailed description.

The CTD of Gemin2 is a helical bundle consisting mainly of 7 α-helices (α2-α8) as reported early (23). In the new structure, yet, there is also a 3_10_-helix after α8 (Figures 1B and 2C-D), which is a short, but conserved structure (Figure 3). The QxDL (residues 272-275) motif is highly conserved sequence. Q272 and D274 form salt bridges with R235 from α7. Interestingly, these three residues are 100% conserved among the orthologues of various eukaryotic species (Figure 3). In addition, Q272 also hydrogen bonds with the main chain amide nitrogen from F270 at the C-terminal end of α8 to maintain the 3_10_-helix conformation. Leu275 faces the interface between α7-α8. The hydrophobicity of this residue is conserved (Figure 3). D277 form salt bridges with R245 from α7. These salt bridges are also highly conserved (Figure 3). These further stabilize the 3_10_-helix interaction with its neighboring α7-8. Recently, the importance of the QxDL motif was signified in the Gemin2 orthologue protein Brr1 in *S. cerevisiae* by a genetic approach. The truncated *BRR1-CΔ6* allele encoding Brr1(1–335), in which the last hexapeptide <u>Q</u>K<u>DL</u>IE(336–341) was deleted, and the Brr1 double mutant Q336A/D338A failed to act as the wild-type Brr1 in complementing *brr1Δ/SMF-F33A* and *brr1Δ/SME-K83A*(51).

**Figure 3.**
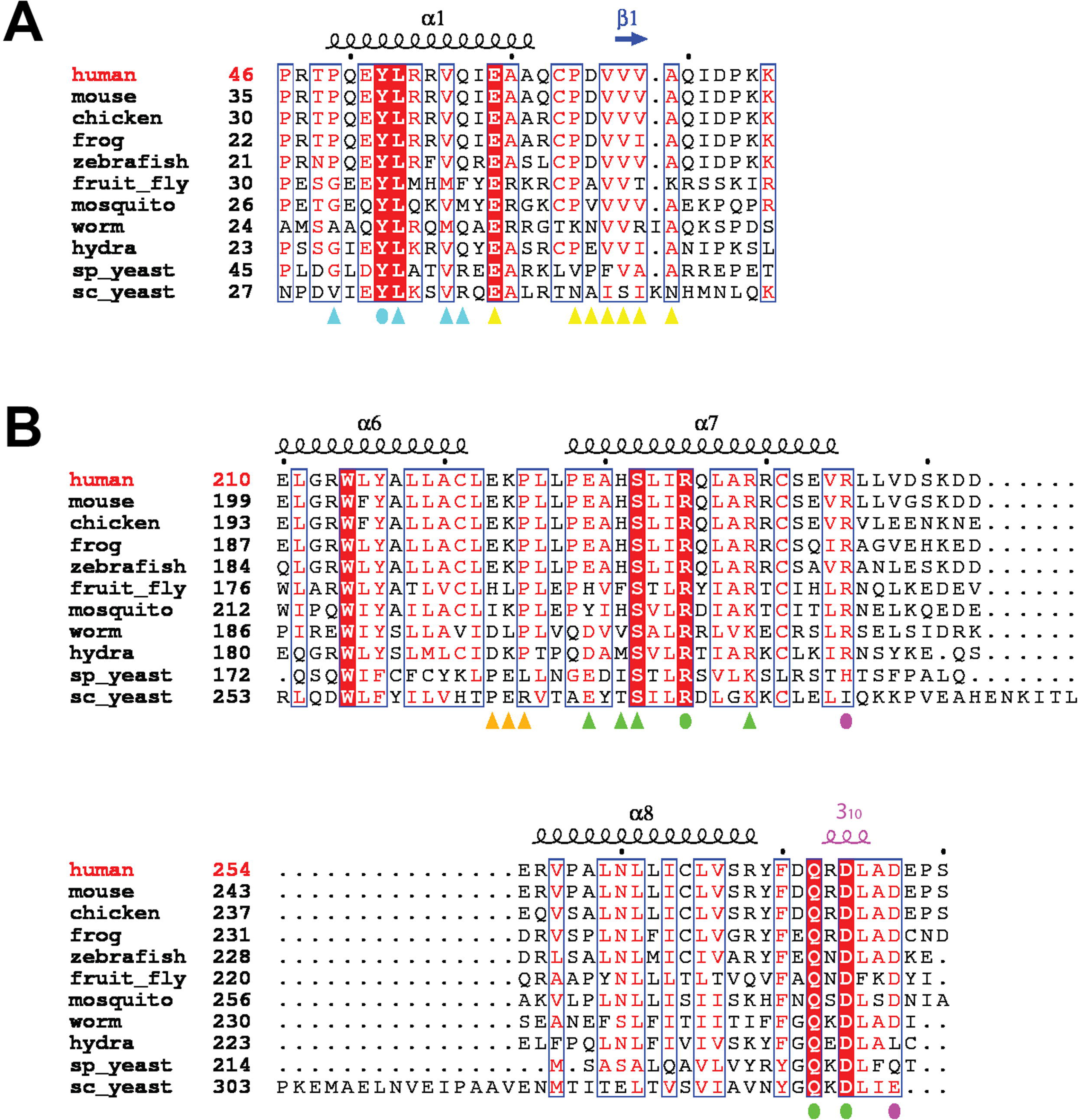
The residues of Gemin2 orthologues contacting Sm proteins in diverse organisms are conserved. Multiple sequence alignment was performed by Clustal Omega (49) and the figure was prepared by ESPript (50). (A) Gemin2’s NTD. (B) Part of Gemin2’s CTD interacting with SmD1/D2. Human, *Homo sapiens* (accession number NP_003607); mouse, *Mus musculus* (NP_079932); chicken, *Gallus gallus* (NP_989530); frog, *Xenopus laevis* (NP_001087945.2); zebrafish, *Danio rerio* (NP_001017608); fruit_fly, *Drosophia melanogaster* (NP_649092.1); mosquito, *Anopheles gambiae* (XP_001688660.1); worm, *Caenorhabditis elegans* (NP_001022847.1); hydra, *Hydra magnipapillata* (XP_002160350); sp_yeast, *Schizosaccharomyces pombe* (CAB88094); sc_yeast, *Saccharomyces cerevisiae* (NP_015382.1). Absolute identical residues (100%) are in white font with red background. Highly conserved residues (over 70%) are in red font and boxed in blue. The secondary structure of human Gemin2 is shown on the top. Cyan, yellow, orange and green triangles indicate the residues contacting SmE, SmF, SmD1 and SmD2 respectively. Cyan circle indicates the residue contacting both SmE and SmF. Green circles indicate the residues contacting each other as well as SmD2. Pink circles indicate the residues contacting each other.

The interactions of Gemin2’s CTD with SmD1 are the same as described in our previous study (23). Briefly, the N-terminal parts of α5 and α7 and the loop between α6-α7 of Gemin2 make contacts with β2 near loop 2 and β3-β4 near loop4 of SmD1, and most of the contacts are hydrogen bonds (Figure 2E). However, particular attention should be paid to the interaction pattern: the interacting atoms of Gemin2 are mostly from the main chain. This may explain why the sequence of the loop between α6-α7 in Gemin2 is not conserved but its length indeed is, indicating a less flexibility on this loop (Figure 3). On the other hand, the interacting residues in SmD1 are mostly located near loops 2 and 4, where variations are possible (Supplementary Figure S1).

The interactions of Gemin2’s CTD with SmD2, however, are more than described previously (23). The main interacting parts are α7 and 3_10_-helix from Gemin2 and β3-β4 from SmD2 (Figure 2F). From Gemin2’s α7, R235 and R239 make salt bridges with D93.SmD2, and H231 makes hydrogen bond with R94.SmD2, as described previously (23). In addition, S232 hydrogen bonds the amide nitrogen of D93.SmD2 and the side chain of E229 hydrogen bonds with the side chain of W74.SmD2. From Gemin2’s 3_10_-helix, Q272 forms hydrogen bond with the amide nitrogen of Y95.SmD2, and R273 forms a hydrogen bond network with the side chains of N69 and Y95 from SmD2. Overall, the interactions between Gemin2’s CTD and SmD2 are mostly polar and are intensive. Because of the above described features, the interacting surface of Gemin2 to SmD1/D2 is highly rigid and can be viewed as a solid rock surface.

In metazoans, the assembly of U7 snRNP cores take the same way as spliceosomal snRNP cores do, and are also mediated by the SMN complex (5,6,13). U7 Sm cores share five Sm proteins with spliceosomal snRNP cores, but have Lsm10/11 heterodimer in the place of SmD1/D2. The sequence alignments of human Lsm10 and Lsm11 versus SmD1 and SmD2 from various species reveal that majority of the residues of Lsm10/11 to interact with Gemin2’s CTD are maintained or replaced by similar residues (Supplementary Figure S1). So, we can predict that Gemin2 binds Lsm10/11 and SmF/E/G in a similarly narrow conformation and applies a similar constraint role on U7 snRNAs.

### RNA-dependent widening of 5Sm and RNA selection

RNA binding is the cause of the splay of 5Sm from the 7S intermediate to become the Sm subcore. Although the structure of the Sm subcore has not been experimentally determined, it should have the conformations of both RNA and 5Sm very similar to those in the mature Sm core for the following reasons: (1) the Sm subcore should have the opening between SmD1 and SmG not narrower than the distance between them in the Sm core because SmD3/B would otherwise be unable to join; (2) The opening between SmD1 and SmG should not significantly larger than the distance between them in the Sm core because the circular form of the Sm site RNA in the Sm core would be the conformation with the lowest energy. Therefore the structure of the Sm subcore is modeled by simply removing SmD3/B from the Sm core structure (Figure 1B) and used in the following analysis. Remarkably, the conformation of the Sm site RNA inside Sm subcores and cores (44,45) is unusual because the RNA backbone, which is negatively charged, forms a ring inside a very narrow space with the bases protruding outside to contact Sm proteins (Figures 1B and 4A). There must be very strong repulse among the negative backbone charges. This conformation of RNA would hardly exist in itself. The existence of such a highly tensive ring of the Sm site is only possible when there are very intensive specific interactions between the 9 Sm site bases and at least 5 Sm proteins (for Sm subcore). The recent U1 and U4 crystal structures describe the detailed interactions between the Sm site bases and 7 Sm proteins (44,45). These interactions form the primary structural basis for snRNA binding specificity in Sm core assembly.

When superimposing SmD2 of the snRNP cores and 7S, there are clearly clashes of SmE and SmG with the Sm site RNA and of SmE with the 3’-SL RNA (Figure 4A), indicating that the RNA conformation in its initial binding to the 7S is different from that in Sm subcores and cores. To make a Sm site model in the 7S Sm heteropentamer, the first 7 bases of the Sm site in U4 snRNP were individually saved and linked again after individual Sm protein is aligned with the corresponding one in the 7S, followed by a relaxing of conformational constrains. In contrast to a circle seen inside U4 snRNP, the Sm site RNA inside the 7S is an ellipse with two bases (U4 and U5) bulging out at the SmD1-SmG opening (Figure 4B-C). The negative charges at the opposite sides of the minor axis of the ellipsoid Sm site RNA backbone would repulse each other much more than those at the opposite sides of the major axis. In other words, the evenly distributed negative charges on the backbone tend to splay the ellipsoid Sm site RNA into a circle, which provides the transition force to widen 5Sm (Figure 4D).

**Figure 4.**
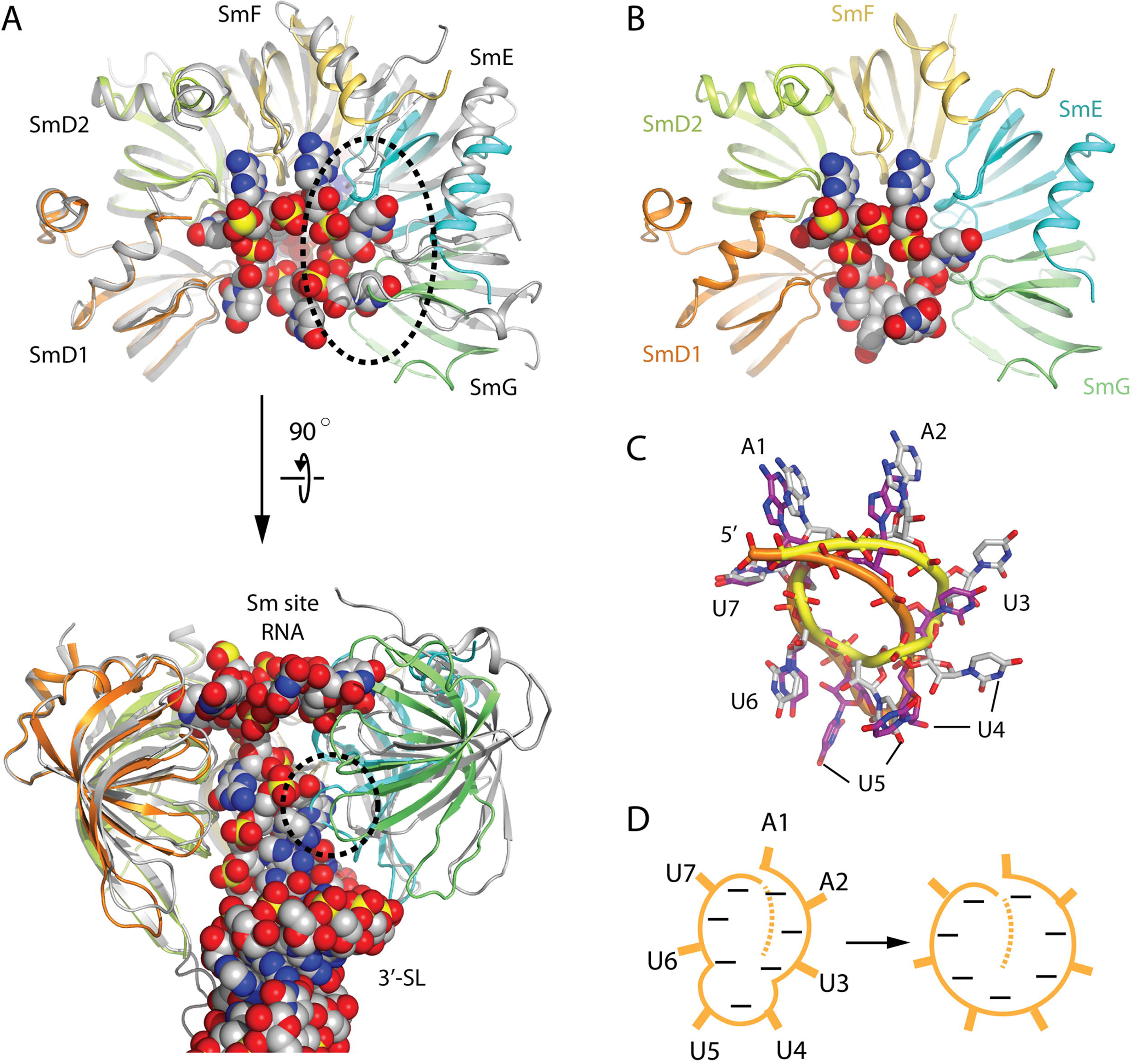
RNA-dependent widening of 5Sm. (A) Comparison of 5Sm in the 7S intermediate (PDB code 5XJL) and the Sm core (represented by U4 snRNP core, PDB code 4WZJ)(45). SmD2 is used for superposition. The five Sm proteins in the 7S are colored the same as in Figure 1B. They are colored in light grey in the Sm core. The collided portions between the RNA in the Sm core and the 5Sm in 7S are indicated by black dashed circles. (B) The Sm site RNA model in the 7S. RNA in (A) and (B) is showed in sphere. Oxygen, nitrogen and phosphor in RNA are colored in red, blue and yellow respectively. (C) Comparison of the Sm site RNA model in the 7S (carbon is colored in pink and phosphate backbone in orange) with the Sm site in the Sm core (carbon is colored in light grey and phosphate backbone in yellow). Numbering starts from the first nucleotide in the Sm site. (D) The ellipsoid Sm site RNA (in the 7S) tends to expand into a circle.

A nonameric Sm site RNA oligonucleotide (9nt), AAUUUUUGA, is able to form a stable Sm subcore with SmD1/D2 and SmF/E/G alone (15), however it cannot bind to the 7S (43). The formation of the Sm subcore from SmD1/D2 and SmF/E/G alone should have a transient 5Sm complex first before the binding of 9nt. Although we don’t know whether there is a narrow conformation of the 5Sm complex, if there is, the conformation of 5Sm would be quite flexible to be able to allow the binding of 9nt and an easy splay of 5Sm. The intensive specific interactions between 9nt and 5Sm seem sufficient to provide the energy to keep the Sm site RNA stably binding to 5Sm and very reasonably in a circular conformation as it is in the mature Sm core. However, in the 7S, because the binding of Gemin2 from the periphery of 5Sm constricts them in a stable, narrow conformation, the binding energy provided solely from the interactions between 9nt and 5Sm seem not enough for 9nt RNA to bind to 5Sm in the 7S. We can image that a very transient complex would have the 9nt Sm site RNA bound to 5Sm but staying in a highly tensive ellipsoid shape and therefore the RNA would be immediately repealed out of 5Sm. To bind to the 7S, there must be an additional RNA at the 3’ side of the Sm site, which can provide additional interactions with 5Sm and additional energy. This explains why a U4 RNA derivative keeping the 5’ side of the Sm site intact but deleting the 3’ side is unable to bind to the 7S (43).

The best candidate at the 3’ side of the Sm site is a sequence-independent stem-loop, which has a minimal 5-6 base pairs, immediately following the Sm site. These features together with the Sm site were termed as snRNP code and are characteristic features of snRNAs (11). A detailed analysis of RNA-Sm protein interactions in U4 snRNP (PDB code 4WZJ) explains why it is the case. There are many positively charged residues on the inner side of 5Sm to interact with both strands of the 3’-SL (Figure 5A and F). From the downward strand of 3’-SL, R47 and N49 from loop 2 in SmD2 interact with the phosphate backbone at the 1^st^ and 3^rd^ base pairs (Figure 5C and F), R66 from β5 and K48 from loop 2 in SmD1 contact the phosphates at the 2^nd^ and 3^rd^ base pairs (Figure 5B and F). Going further down, there are a group of positively charged residues, K79, K82 and K85 in loop 4 from SmD2, contacting the phosphates at the 5^th^-6^th^ base pairs (Figure 5C and F). In addition, the RNA upward strand also contacts Sm proteins. R50 in loop 4 of SmD1 forms salt bridges with the backbone between the 8^th^ and 9^th^ base pairs (Figure 5B and F). K84 in loop 4 of SmD2 forms salt bridges with the backbone between the 10^th^ and 11^th^ base pairs (Figure 5C and F). K22 at β1 near loop 2 of SmF contacts the phosphate at 3’-end of RNA (Figure 5D and F). SmE makes more polar contacts with the RNA upward strand (Figure 5E and F): K67 at loop 4 contacts the phosphate between the 2^nd^ and 3^rd^ base pairs, N40 at loop 2 contacts the phosphate between the 1^st^ and 2^nd^ base pairs, Q38 at loop 2 contacts the phosphate at the free RNA 3’-end, and H65 at loop 4 hydrogen bonds the 2’ oxygen of the last ribose. These charged interactions provide additional binding energy for RNA to bind into the central channel of 5Sm and for the Sm site to splay into a circle, which widens the opening between SmD1 and SmG. Almost all these interactions are between the RNA backbone and the Sm proteins. This explains the sequence independence of the 3’-SL in RNA binding (11). The size of 3’-SL had been studied on U7 snRNA to test its influence on U7 Sm core assembly and it was found that less than 5 base pairs in the 3’-SL impair U7 Sm core assembly (12). It is consistent with the fact that majority of the interactions of 3’-SL with Sm proteins are located within the first 5 base pairs. As there are substantial interactions between the upward backbone of the first 3 base pairs of the 3’-SL and SmE/F, especially SmE, a SL immediately adjacent to the 3’ end of the Sm site would be preferred for Sm core assembly. This also gives a good explanation of an *in vivo* selection experimental result (52): a library of 20 random nucleotides (N20) inserted into the position between two SLs of a U1 snRNA derivative, replacing the original Sm site, was injected into oocytes, and the assembled Sm cores in the nucleus were enriched and selected by 12 rounds of Sm-specific Y12 antibody precipitation, PCR amplification and injection. The selected RNAs assembled into Sm cores all contain AAUUUUUGG at the 3’ end of N20, immediately adjacent to a 3’-SL.

**Figure 5.**
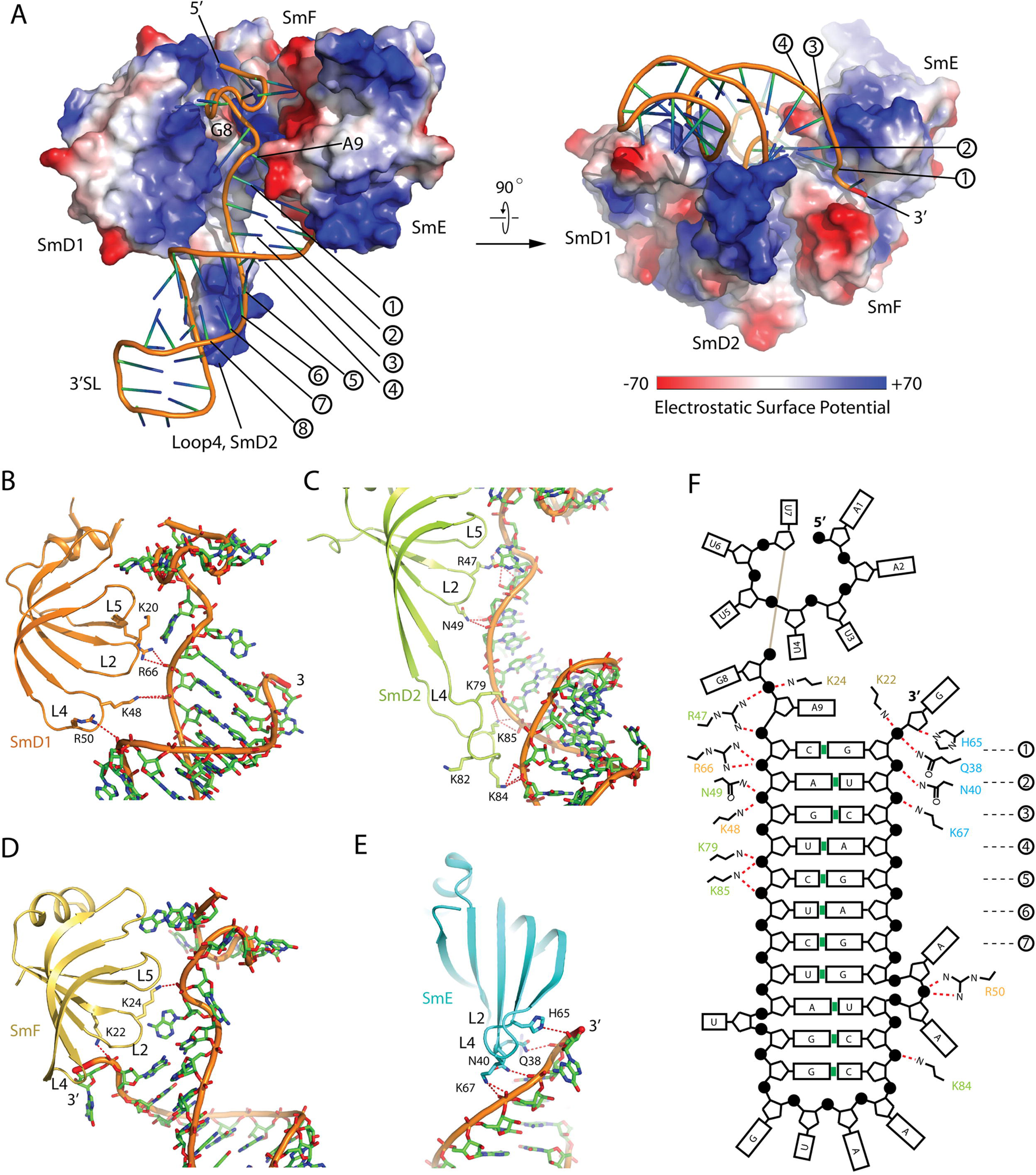
Interactions between Sm proteins and RNA 3’-SL are critical for RNA binding selectivity. (A) Interactions of the snRNP code with SmD1/D2/F/E in the Sm core (PDB code 4WZJ) (45). Because of its little contact with RNA 3’-SL, SmG is not shown for clarify. Sm proteins are shown in surface view with electrostatic surface potential. RNA is shown in cartoon. The base pairs immediately after the Sm site RNA are numbered. (B-E) Detailed interactions between each Sm protein and RNA 3’-SL. (F) Schematic summary of the interactions between Sm proteins and RNA 3’-SL.

For biochemical characterization, a single-stranded RNA 3’-downstream of the Sm site seems sufficient to allow RNA to bind into the binding site and to widen the SmD1-SmG opening (43). An *in vivo* experiment in which injection of a ‘de-stem’ U4 RNA (equivalent to the Sm site plus a 3’ single strand) into oocytes could be assembled into the Sm core albeit with low efficiency also demonstrates this (11). A single-stranded RNA would keep most of the polar interactions with SmD1 and SmD2, as the downward strand of the SL does. But SL structure is more efficient, as it not only provides additional interactions to SmF and SmE, but also offers a more rigid double-stranded 3’-RNA for better interactions with Sm proteins. This feature explains why snRNAs having a 3’-SL immediately downstream of the Sm site dominate the Sm core assembly inside cells (18).

The interactions between Sm proteins and 3’-SL RNA are also highly conserved among eukaryotes. For example, R47 and N49 in SmD2, R66 in SmD1 and K22 in SmF are 100% conserved (Supplementary Figure S1). K48 and R50 in SmD1 and K79 and K85 are highly conserved, the changes being mostly from K to R or R to K (Supplementary Figure S1). A special case is SmD2 in *Saccharomyces cerevisiae* in which its loop 4 is shorter than those of the other organisms and loses several RNA-interacting residues (Supplementary Figure S1). However, loop 4 of SmD1 in *S. cerevisiae* is 27 residues longer than those of other organisms and it contains several fully or partially positively charged residues, K, N and Q (Supplementary Figure S1), which would potentially interact with 3’SL and compensate for the loss of interaction of loop 4 of SmD2 with RNA.

As Lsm10/11 replaces SmD1/D2 in the assembly of U7 snRNP core(5,6), comparison of the sequences of Lsm10 versus SmD1 and Lsm11 versus SmD2 also gives rise to a high conservation of 3’-SL-interacting residues in Lsm10/11 (Supplementary Figure S1). K20, K48 and R66 in SmD1 have R34, R62 and R80 in their equivalent positions in Lsm10. R47 and K82 in SmD2 have the same residues in Lsm11 in their equivalent positions. This high conservation explains similar interactions between Lsm proteins and 3’-SL and a similar requirement of 3’-SL of U7 snRNA in U7 Sm core assembly.

### Conformational changes of 5Sm upon RNA binding

In the previous analysis by comparison of 5Sm in the 7S and U1 snRNP, it was suggested that the interfaces within each of the Sm subcomplexes, SmD1/D2 and SmF/E/G, are rigid, but the SmD2-SmF interface is widened in U1 snRNP (23). However, this suggestion cannot explain the negative cooperativity between Gemin2 and RNA in binding to 5Sm, because the segment of Gemin2 corresponding to the widening SmD2-SmF interface is only the flexible connecting loop between its NTD and CTD (Figure 1B). In the previous analysis, the entire structures of Sm proteins were used for superimposition (23), which might shield potential subtle changes of their N-terminal helices and the following β sheets. To make a better comparison, in this study, 48 residues of each Sm protein, which are from the β sheet and are less varied, are used (Figure 6A). In addition, the recent structures of U1 and U4 snRNP cores, which have the highest resolutions of 3.3 and 3.6 Å respectively, are chose for the comparison (PDB codes 4PJO and 4WZJ) (44,45). Because there are 4 U1 snRNP molecules and 12 U4 snRNP molecules in one asymmetric unit (ASU) respectively, the similarity of the sixteen 5Sm molecules was analyzed at first. Superposing the same Sm protein from any two of the four U1 snRNP molecules, the root mean square deviations (RMSDs) of the 192 main chain atoms of other Sm proteins are very small (Supplementary Figure S2A), indicating almost identical conformation of the four 5Sm in each ASU. Superposition of the twelve U4 snRNP core molecules also produces a similar result. Then a representative 5Sm in a U1 snRNP core is compared with one from U4 snRNP core. When each pair of the corresponding Sm proteins is superposed, the neighboring Sm protein pairs are also superposed well, as indicated by low RMSD values (not larger than 0.83 Å for 192 main chain atoms, although they are a little larger than those of U1 or U4 core molecules in ASU) (Supplementary Figure S2, B-D). This indicates that the conformations of the sixteen 5Sm molecules in U1 and U4 snRNP cores are very similar and any of them is proper to represent the conformation of 5Sm in the Sm subcore and core.

**Figure 6.**
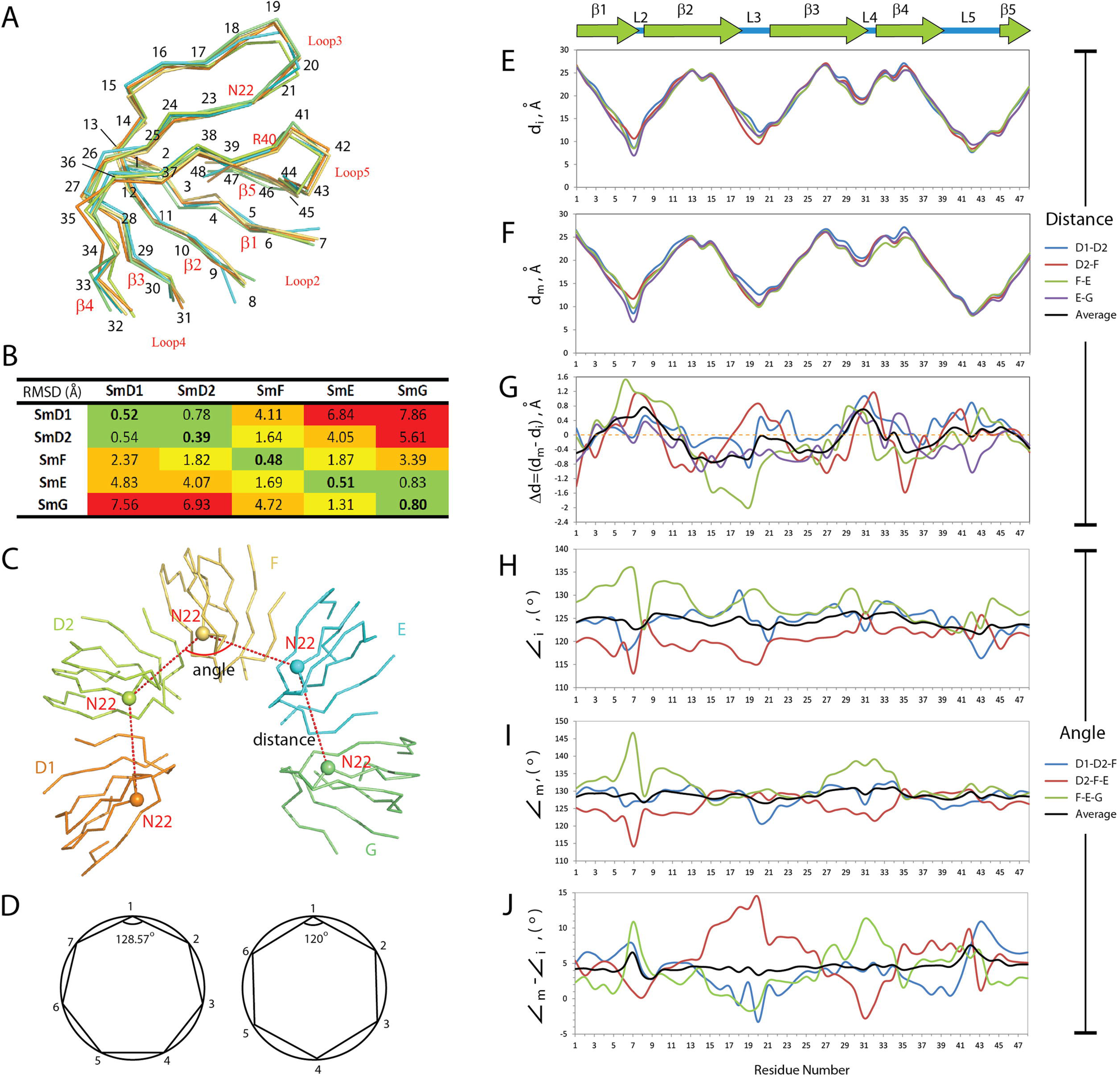
Conformational changes of 5Sm from the 7S intermediate state to the assembled Sm subcore/core. (A) The Sm fold defined by the 48 residues of each Sm protein used for superposition and analysis. Numbering residues in this system is shown at each position. (B) Position differences of each pair of Sm proteins in the 7S and the Sm core (U1 snRNP core, PDB code 4PJO)(44) when one pair is superposed using the 48-residue Sm fold system. (C) Definition of the distance and angle used in analysis. (D) Standard inner angles for regular heptagon and hexagon. (E-J) Conformational changes of 5Sm from the 7S to the Sm subcore/core characterized by the above-defined distances and angles. Distances between each pair of positions in adjacent Sm proteins in the 7S intermediate, d_i_ (E), and in the mature Sm core, d_m_ (F), are plotted against the 48 positions. (G) Distance changes from the 7S to the Sm core, Δd = d_m_-d_i_, are plotted against the 48 positions. Angles of three adjacent Sm proteins in the 7S, ∠_i_ (H), and in the Sm core, ∠_m_ (I), are plotted against the 48 positions. (J) Angle changes from the 7S to the Sm core, ∠_m_-∠_i_, are plotted against the 48 positions. The secondary structures of the 48 positions are shown on the top of the plots.

5Sm in the 7S and one in a U1 snRNP are used for comparison. To quantify the variations of each pair of identical Sm proteins, the RMSDs of the above-mentioned 48 residues of all 5 pairs of Sm proteins were calculated when each pair is superposed. While the RMSDs of the five Sm proteins are relatively small if superposing individually (0.52, 0.39, 0.48, 0.51 and 0.80 Å for the 192 main chain atoms of D1, D2, F, E and G respectively), the RMSDs of their adjacent Sm proteins increase, albeit to different extent (Figure 6B), indicating splay between each adjacent Sm protein pair in 5Sm. This is distinct from our previous analysis, in which only the splay between SmD2 and SmF is noticed (23). Specifically, the RMSD increases of SmD1 (from 0.52 to 0.54 Å when SmD2 is superposed) and SmD2 (from 0.39 to 0.78 Å when SmD1 is superposed) are relatively small (Figure 6B), indicating a subtle splay between SmD1 and SmD2. However, the RMSD increases of SmD2 (from 0.39 to 1.82 Å when SmF is superposed) and SmF (from 0.48 to 1.64 Å when SmD2 is superposed) are large (Figure 6B), indicating a large splay between SmD2 and SmF. This conclusion is consistent with our previous study (23). However, the splay between SmF and SmE is also large as indicated by the large RMSD increases of SmF (from 0.48 to 1.69 Å when SmE is superposed) and SmE (from 0.51 to 1.87 Å when SmF is superposed) (Figure 6B). Finally, the RMSD increases of SmE (from 0.51 to 1.31 Å when SmG is superposed) and SmG (from 0.80 to 0.83 Å when SmE is superposed) are also perceivable (Figure 6B), indicating conformational changes between SmE and SmG. To summarize, although there are overall increased conformational changes between all the neighboring Sm proteins, the conformational changes of SmD2-SmF and SmF-SmE are more substantial.

To make more detailed analysis of the conformational changes between Sm proteins, two more parameters are used (Figure 6C): the distance between the Cα of the 48 residues of each pair of adjacent Sm proteins, d_n_(Sm_1_-Sm_2_), and the angle of the Cα of the 48 resides from 3 adjacent Sm proteins, ∠_n_(Sm_1_-Sm_2_-Sm_3_). For each adjacent Sm pair, either in the 7S or in U1 snRNP, distances are small at the loop regions, which face the central RNA-binding channel, and large at the perimeter (Figure 6, E-F). The distance changes, Δd_n_ (= d_m_-d_i_), (m, mature Sm core, and i, intermediate 7S), are mainly positive at each loop and negative at the perimeter (Figure 6G), consistent with the overall splaying of the horseshoe-shaped 5Sm. Large distance increases occur at two loop regions, loops 2 and 4, both of which are located at the lower part of 5Sm interacting with 3’-SL RNA. The average Δd_n_ are between −0.8 and 0.8 Å, indicating a small adjustment upon snRNA binding to 5Sm. However, when individual adjacent Sm pairs are examined, the changes are not equal at their corresponding positions. The distance changes are relatively small for SmD1-SmD2 and SmE-SmG, while larger changes occur for SmD2-SmF and SmF-SmE. This is consistent with the insertions of larger bases between the adjacent Sm proteins: A1 into between SmD2 and SmF, and A2 and A9 into between SmF and SmE.

The average angle is 124.27° for the 7S and 128.80° for U1 snRNP (Figure 6H and I). Considering the inner angles of 120° in a regular hexagon and of 128.57° in a regular heptagon (Figure 6D), the angle for U1 snRNP is very close to the standard value of a heptamer, but the Sm pentamer in the 7S occupies 5 positions of an approximately 6.5-Sm-forming ring. Interestingly, the average angles of D1–D2-F, D2-F-E, and F-E-G are not similar (Figure 6H and I). They are 124.46°, 120.45° and 127.92° in the 7S, and 128.49°, 126.13° and 131.79° in U1 snRNP. This indicates that the angle from D2-F-E in the 7S is smaller than those of other adjacent Sm proteins and is close to the standard angle of a hexamer. In addition, although the average angle change, Δ∠ (∠_m_ - ∠_i_), is 4.53°, the average angle change for each set of adjacent Sm proteins varied (Figure 6J). They are 4.03°, 5.68° and 3.87° for D1–D2-F, D2-F-E and F-E-G. This indicates that the angle from D2-F-E is widened more upon RNA binding and Sm subcore formation.

In addition, the angle changes over the 48 positions vary very much (Figure 6J). At loop 3, Δ∠_18-22_ (D2-F-E) increases dramatically, reaching a value of about 15°, about three-fold the average change. This is caused by the insertions of two adenines into D2-F and F-E respectively, consistent with the distance changes at this location stated above. Correspondingly, the Δ∠_18-22_ (D1–D2-F) and Δ∠_18-22_ (F-E-G) become negative. At loops 2 and 4, Δ∠ (F-E-G) are larger than the average while Δ∠ (D2-F-E) are close to zero and even becomes negative. This is caused by the 3’-end of RNA passing between loops 4 of SmF and SmE (Figure 5A). In summary, the conformations of individual Sm proteins and the interfaces between adjacent Sm proteins are changed, but not evenly, which are affected by the local environment of the bound RNA.

### Structural basis for Gemin2’s release

The binding of snRNA to 5Sm in the 7S complex triggers the release of Gemin2 (43). The widening of 5Sm accompanies the splay of the Sm site RNA is the major reason. As analyzed above, the changes occur all over the Sm proteins, although most of them are quite subtle. Because there is no apparent constraint exerted on the interface between SmD2 and SmF by Gemin2, the subtle conformational changes of the interactions between Gemin2’s NTD and SmF/E and between Gemin2’s CTD and SmD1/D2 are the major source for Gemin2’s release.

Superposing SmE or SmF can generate a slight twist and location shift. When superposing SmE based on the 48 residues, for example, the distance between the Gemin2-interacting residues are slightly altered (Supplementary Figure S3A). The Cα-Cα distance between I18 and E52 decreases 0.4 Å, while that between L25 and F50 increases only 0.2 Å. When superposing SmF on the basis of the 48 residues, a rigid-body shift of helix 1 toward periphery is observed (Supplementary Figure S3B). The Cα-Cα distance between N12 and G38 increases 1.7 Å, while that between N12 and V34 decreases 1.1 Å. The distance between helix 1 of SmE and β2 of SmF, the two structural parts marking the edges of the Gemin2’s NTD-SmE/F interaction set, is reduced (Figure 7A-B). The Cα-Cα distance between I18.SmE and G38.SmF decreases 2.3 Å, from 22.9 to 20.6 Å. It is just about 10% reduction. How can such a small change of the SmE/F make Gemin2’s NTD get away? One might think that such a small change can be accommodated by subtle adjustment of protein as we commonly consider that proteins are usually plastic. However, the key point here is that Gemin2’s NTD is like two sticks linked by a joint as described above. When helix 1 of SmE moves towards SmF, the "ridges into grooves" interaction between the N-terminal α1 of Gemin2 and helix 1 of SmE makes the entire Gemin2’s α1 move towards SmF too, the extra 2.3 Å spatial tension needs the 3-residue linker to release (Figure 7A). There are only two ways to do so: (1) popping the linker inward to the central RNA-binding channel; (2) popping the linker outward to the periphery (Figure 7A and C). In case (1), the linker moves in the direction opposite to that of its interacting residues K8 and N12 of SmF, and the hydrogen bond network breaks. In addition, as the connection between α1 and the linker is covalent, the C-terminal part of α1 would also move in the direction of the linker. This would increase the distance between Y52, E59 in Gemin2 and N6 in SmF, and the hydrogen bonds between them would break too. In this way, multiple hydrogen bonds would be lost (Figure 7C). In case (2), the linker moves in the same direction as that of its interacting residues K8 and N12 of SmF, the hydrogen bond interactions mentioned above would be kept. However, the extra 2.3 Å length tension would be relayed to β1 of Gemin2. As the distance between two adjacent Cα atoms in a β strand is about 3.8 Å, the extra length of 2.3 Å is too short to shift the register of the interacting residues by one to make hydrogen bonding again. Instead, the distance between adjacent hydrogen bonds from antiparallel β strands is about 2.4 Å, the 2.3 Å relay would actually move H-donor nitrogen on Gemin2’s β1 to face H-donor nitrogen from SmF’s β2. All the hydrogen bonds between this pair of antiparallel strands would not only be lost, but the elements on the strands otherwise hydrogen bonding would be in a repulse mode. In this way, multiple hydrogen bonds would lose too (Figure 7C). In either way, the "stick-joint-stick" of Gemin2’s NTD would no longer fit into the binding sets of SmF/E after the splay of 5Sm.

**Figure 7.**
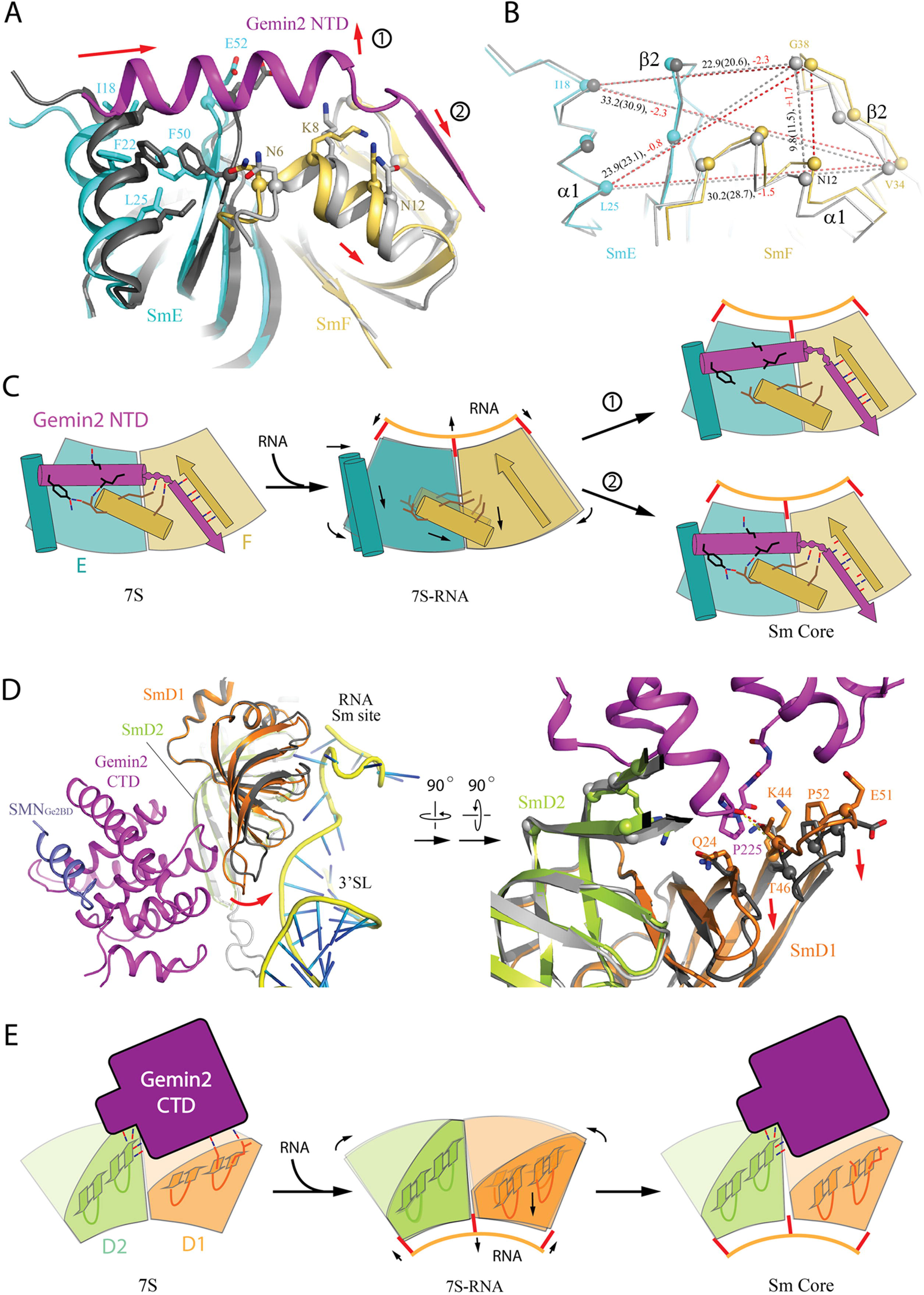
Structural basis for Gemin2’s release coupled with RNA-dependent 5Sm conformational changes. (A) Potential changes of the interactions between Gemin2’s NTD and SmE/F upon Sm core assembly. SmFs in the 7S and Sm core are superposed based on the 48 positions defined previously. Gemin2’s α1 would move towards SmF following the N-terminal helix of SmE as indicated by the red arrow. The second half of Gemin2’s NTD would have two types of possible movements as indicated by red arrows labeled with ① and ②. (B) Distance changes between the indicated positions on Sm proteins from the 7S to the Sm subcore/core. SmEs in the 7S and Sm core are superimposed based on the 48 positions defined early. Distances (Å) in the 7S and in the Sm core are shown in black with the latter in brackets, and distance changes are in red. (C) Schematic model of Gemin2’s NTD reducing contacts with SmE/F. (D) Changes of the interactions between Gemin2’s CTD and SmD1/D2 upon Sm core assembly. Superimposition of SmD2s of the 7S and Sm core reveals a flipping of SmD1’s loops 2 and 4 towards RNA 3’-SL (red arrows), which causes Gemin2’s CTD to lose most contacts with SmD1. (E) Schematic model of Gemin2’s CTD reducing contacts with SmD1/D2 upon Sm core assembly. Hydrogen bonds are shown in red (H-acceptor) and blue (H-donor) bar pairs in schematic models (C) and (E).

The superposition of SmD2 does not make many shifts of the interacting residues from SmD2, as the Cα atoms only move within short distances of 0.1-0.2 Å (Figure 7D). However, loops 2 and 4 in SmD1 move closer to 3’-SL RNA (Figure 7D), because of the positively charged residues at these loops making polar contacts with the RNA backbone. As the interacting residues of SmD1 with Gemin2’s CTD are exclusively located in loops 2 and 4 (Figure 2E), the flipping of the loops of SmD1 towards the bound RNA increases the distance between the interacting surfaces of SmD1 and Gemin2’s CTD (Figure 7D). And as the SmD1/D2-interacting surface of Gemin2’s CTD is quite rigid like a rock surface, it would hardly make a plastic adjustment and therefore, the contact between Gemin2’s CTD and SmD1 would be lost (Figure 7D-E).

The widening of 5Sm causes part of SmF/E to lose contact with Gemin2’s NTD and SmD1 with Gemin2’s CTD. By an approximate estimation, about 8-10 hydrogen bonds are lost (about 4-6 between SmF/E and Gemin2’s NTD and 4 between SmD1 and Gemin2’s CTD) (compare Figures 2 and 7). Because the nonameric Sm site with additional 3’-single strand or 3’-SL can bind and widen 5Sm with Gemin2 bound but the Sm site alone cannot (43), the additional interactions between the 3’-RNA and 5Sm can be estimated to compensate for the breaking of hydrogen bonds between Gemin2 and 5Sm. There are about 7-8 polar interactions between the 3’-downstream strand and 5Sm and about 5 between the 3’-upstream strand and 5Sm (Figure 5). Therefore, there are about 12-13 polar interactions for 3’-SL and about 7-8 for 3’-single strand. If one polar interaction is estimated equal to one hydrogen bond, the forming interactions and the breaking ones are at a similar level, and therefore RNA containing the snRNP code or the Sm site plus a 3’-single strand would not cause Gemin2 to release easily. The RNA and Gemin2 would allosterically compete for binding 5Sm, and the 7S and Sm subcore would coexist in equilibrium. This analysis is consistent with our previous observations and provides an explanation at the structural level for them (43).

Our previous study showed that the 7S complex, incubated with both RNA containing the snRNP code and SmD3/B, can completely release Gemin2 and assemble the Sm core (43). This indicated that the joining of SmD3/B into the Sm subcore facilitates Gemin2’s release. Checking the interaction between SmD3/B and the Sm subcore (Figure 1B) provides us the rationale. As analyzed in detail in previous reports of U1 and U4 Sm core structures (1,44,45), there are intensive interactions between SmD3/B and RNA (two Us in the Sm site), as well as between SmD3/B and 5Sm (SmD1 and SmG). The formation of the Sm core from the subcore gains net, large binding energy and should be an irreversible reaction, which drives the release of Gemin2 to a completion.

In summary, in the 7S intermediate state of Sm core assembly, Gemin2 constrains 5Sm in a narrow conformation mostly by its NTD and CTD, which makes the splay of a nonameric Sm site RNA oligonucleotide hard to achieve unless a 3’-SL immediately follows the Sm site RNA, therefore enhancing RNA specificity. The splay of the Sm site causes the conformational change of 5Sm, including splay of 5Sm and the loops of SmD1 moving towards 3’-SL RNA, which reduces the affinity of both the NTD and CTD of Gemin2 to 5Sm and causes Gemin2 to release. As the binding energy provided by RNA containing the snRNP code and that by Gemin2 are at a similar level, the 7S and Sm subcore battle for formation and coexist in equilibrium. The formation of the Sm subcore, even to a relatively low degree, gives a chance for SmD3/B to bind, which further stabilizes the Sm subcore conformation and moves Sm core assembly to a completion. From the above structural analysis, this mechanism should be conserved in all eukaryotes and apply to all kinds of Sm core assembly mediated by the SMN complex, including canonical spliceosomal snRNP cores and special U7 snRNP cores.

### Implications in the evolution of the chaperoning machinery for Sm core assembly

Gemin2 binds 5Sm, identifies cognate RNAs specifically together with the bound 5Sm, and releases Sm core upon finishing the assembly (23,43). Gemin2 is the most conserved protein of the SMN complex (21,22) and interacts with Sm proteins with highly conserved residues (Figure 3). From the evolutionary point of view, Gemin2 is the central protein of the chaperoning machinery for Sm core assembly and could have been the first protein evolved in this pathway. It is consistent with the observation that there is only Brr1, the orthologue of Gemin2 of the 9-component vertebrate SMN complex, is found in one of the simplest eukaryotes, *S. cerevisiae* (53,54). The join of SMN protein might help replace pICln more efficiently as both pICln and SMN proteins are present in *Schizosaccharomyces pombe* (55–58), another single cell eukaryote with more requirement of snRNP core formation as approximately 40% of its genes contain introns (59). Gemin2 plus the N-terminal part of SMN (residues 1-122), which contains the Gemin2 binding domain, binds the 6S complex, which contains 5Sm and pICln, but stalks in the intermediate 8S state, where pICln blocks the entry of RNAs to the binding pocket (17,24). The C-terminal part of the SMN protein may play an important role in releasing pICln from 5Sm. In *Drosophila*, SMN and Gemin2 are sufficient to assemble Sm core efficiently (22). Therefore, Gemin2 and SMN are the minimal assembly machinery in most eukaryotes with less complexity. Gemin5 may have joined in this pathway accidentally by mutation of an ancient WD40 domain-containing protein to gain its RNA binding capability, therefore interfering with Sm core assembly. This hypothesis is supported by the observations that Gemin5 plays multiple roles (60,61). To cope with this accident, cells may have evolved a RNA helicase-containing protein, Gemin3 (34), to take off RNAs from Gemin5, and other proteins, like Gemin4, Gemin6/7/8 and unrip, to aid in transferring Gemin5 to the vicinity of the SMN-Gemin2 assembly line to allow stripping RNAs at site. In vertebrates, the assembly of Sm cores requires ATP hydrolysis (62), which might be the driving force for Gemin3 to take RNA away from Gemin5.

The Sm cores also evolved as the assembly machinery, the SMN complex, became more complex. In simple eukaryotes like yeasts there is no Lsm10/11. But in metazoans, Lsm10/11 appears and assembles into a different Sm core in place of SmD1/D2. This Sm core accepts a slightly different RNA, U7 snRNA, which has a different Sm site but still keeps a 3’ adjacent SL (6,12). More importantly, U7 Sm core assembly can still use the same chaperoning machinery (5,6,13).

### Implications in SMA biology

The illumination of the basic mechanism of Sm core assembly also has important implications in studies of SMA biology. Since Gemin2 plays most of the basic roles in Sm core assembly and there might be a way to assemble Sm cores in any eukaryote without needing the SMN protein, for example, engineering Brr1, the orthologue of Gemin2 in *S. cerevisiae* and the only important chaperone in Sm core assembly in this species, to bind human 5Sm (43). The updated interactions between Gemin2 and 5Sm and the conformational transition during Sm core assembly at the atomic level in this study would help the engineering. In addition, in all SMA patients, there is a *SMN2* gene, which is nearly identical to the *SMN1* gene with the exception of a critical C to T mutant that alters the splicing pattern of its mRNA transcripts. Consequently, the *SMN2* gene produces only small percent of normal SMN protein and large percent of the C-terminal truncated product, SMNΔ7, which is unstable and dysfunctional(63,64). Early study showed that the truncation creates a degradation signal (degron) at the C-terminus of SMNΔ7 and that a mutation, S270A, can inactivate the degron, stabilize SMNΔ7 and rescue viability of SMN-deleted cells (65). This study, together with our previous characterization (43), provide a further reasoning and support for the potential treatment approach based on stabilization of SMNΔ7. Probably, SMNΔ7 subjects both Gemin2 and itself to degradation by binding to Gemin2 via its N-terminal Ge2BD and depletes Gemin2, which causes low efficiency of snRNP core assembly. In contrast, stabilization of SMNΔ7 may exert little influence on Gemin2 level, which possibly maintains the basic Sm core assembly. Following this thought, a stable protein which can bind to Gemin2 in competition with SMNΔ7 might also avoid this passive loss of Gemin2 in cells and act as a new treatment strategy in studies of SMA therapy.

## DATA AVAILABILITY

The improved atomic coordinate of the 7S complex, SmD1/D2/F/E/G/Gemin2/SMN(residues 26-62) has been deposited with the Protein Data Bank under accession code 5XJL.

## SUPPEMENTARY DATA

Supplementary Data are available online.

