## Supplemental Materials for "Structural Insights into Negative Cooperativity between Gemin2 and RNA in Sm-class snRNP Assembly"

Rundong Zhang

It contains

3 supplementary figures (Figures S1-3),

1 supplementary table (Table S1),

and references for them.

A

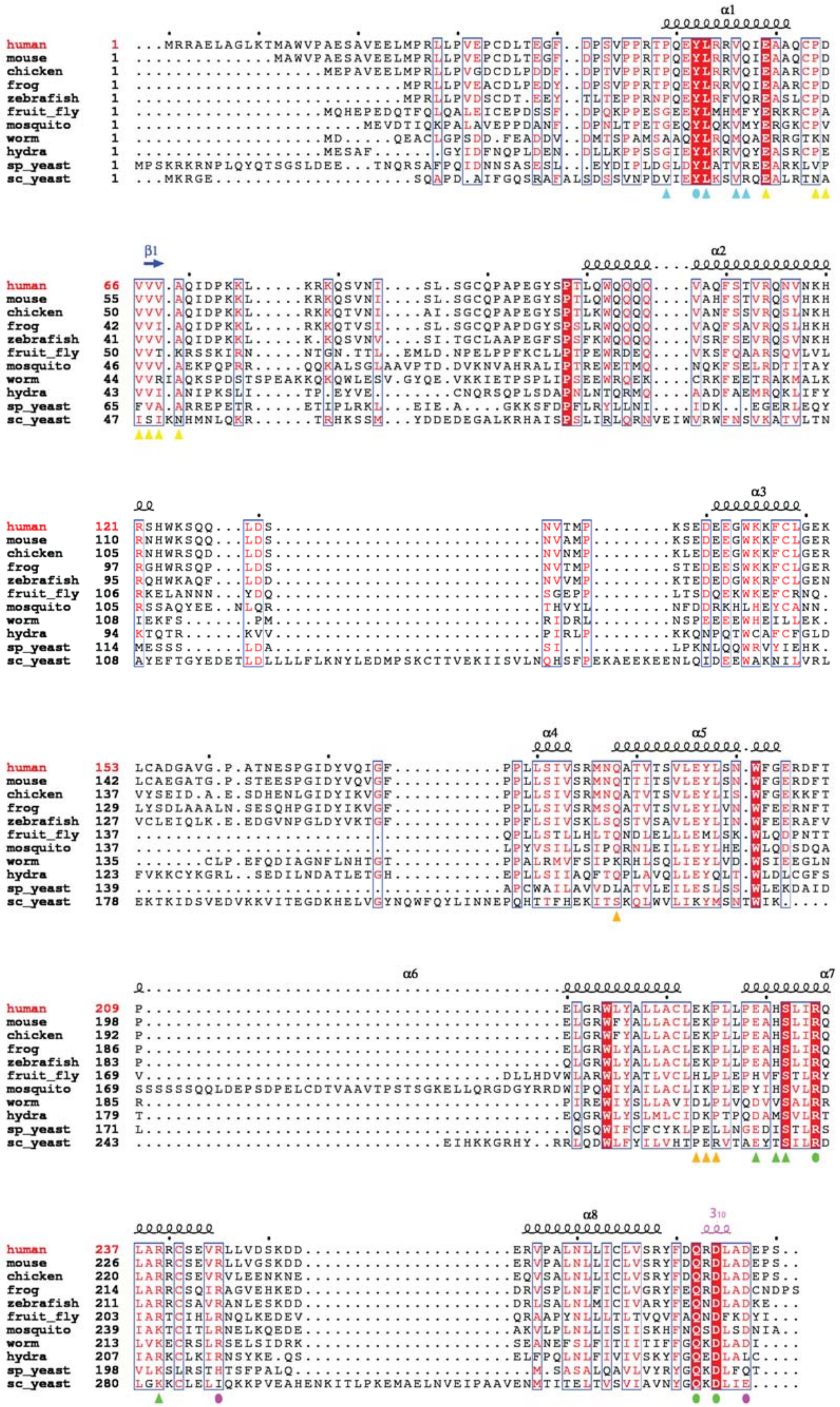
α1
β1
α2
α3
α4
α5
α6
α7
α8
3₁₀
human
mouse
chicken
frog
zebrafish
fruit_fly
mosquito
worm
hydra
sp_yeast
sc_yeast

B

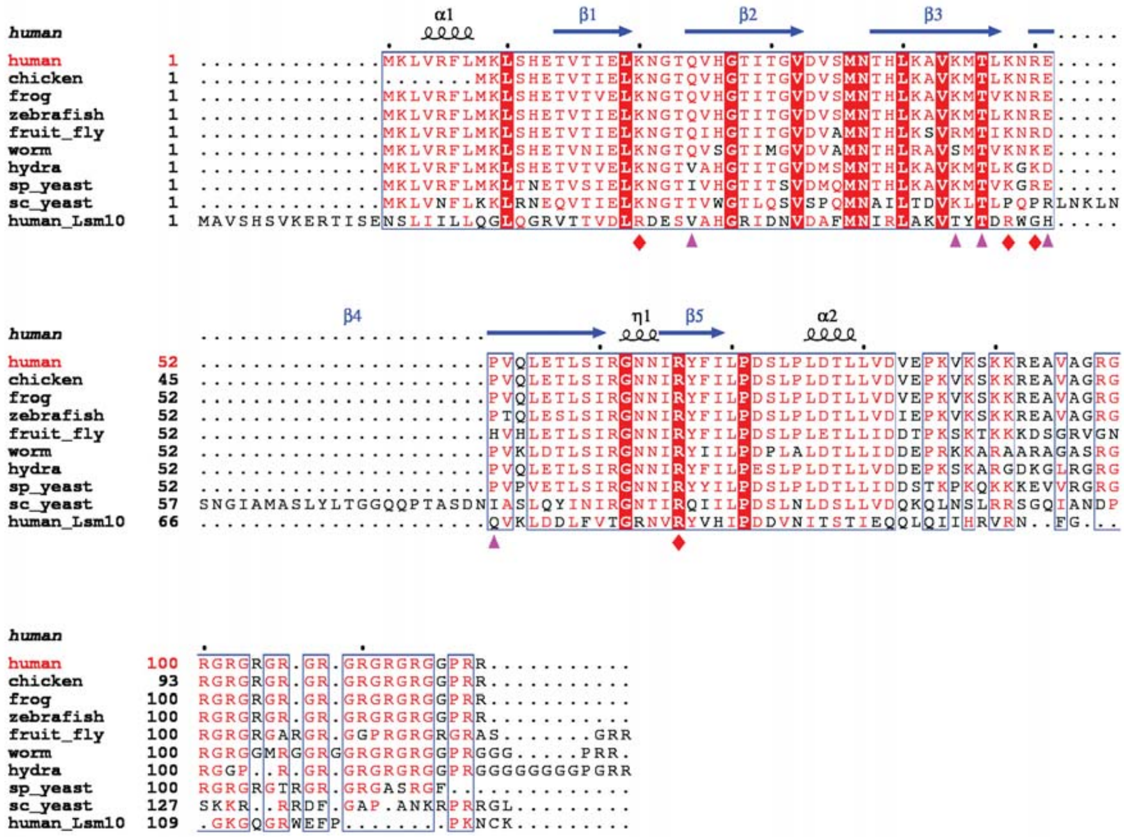

human
α1
β1
β2
β3
human 1 ..............MKLVRFLMKLSHETVTIELKNGTQVHGTITGVDVSMNTHLKAVKMTLKNRE.....
chicken 1 ......................MKLSHETVTIELKNGTQVHGTITGVDVSMNTHLKAVKMTLKNRE.....
frog 1 ..............MKLVRFLMKLSHETVTIELKNGTQVHGTITGVDVSMNTHLKAVKMTVKNRE.....
zebrafish 1 ..............MKLVRFLMKLSHETVTIELKNGTQVHGTITGVDVSMNTHLKAVKMTLKNRE.....
fruit_fly 1 ..............MKLVRFLMKLSHETVTIELKNGTQIHGTITGVDVAMNTHLKSVRMTIKNRD.....
worm 1 ..............MKLVRFLMKLSHETVNIELKNGTQVSGTIMGVDVAMNTHLRAVSMTVKNKE.....
hydra 1 ..............MKLVRFLMKLSHETVTVELKNGTVAHGTITGVDMSMNTHLKAVKMTLKGKD.....
sp_yeast 1 ..............MKLVRFLMKLTNETVSIELKNGTIVHGTITSVDMQMNTHLKAVKMTVKGRE.....
sc_yeast 1 ..............MKLVNFLKKLRNEQVTIELKNGTTVWGTLQSVSPQMNAILTDVKLTLPQPRLNKLN
human_Lsm10 1 MAVSHSVKERTISENSLIILLQGLQGRVTTVDLRDESVAHGRIDNVDAFMNIRLAKVTYTDRWGH.....
human
β4
η1
β5
α2
human 52 ......................PVQLETLSIRGNNIRYFILPDSLPLDTLLVDVEPKVKSKKREAVAGRG
chicken 45 ......................PVQLETLSIRGNNIRYFILPDSLPLDTLLVDVEPKVKSKKREAVAGRG
frog 52 ......................PVQLETLSIRGNNIRYFILPDSLPLDTLLVDVEPKVKSKKREAVAGRG
zebrafish 52 ......................PTQLESLSIRGNNIRYFILPDSLPLDTLLVDIEPKVKSKKREAVAGRG
fruit_fly 52 ......................HVHLETLSIRGNNIRYFILPDSLPLETLLIDDTPKSKTKKKDSGRVGN
worm 52 ......................PVKLDTLSIRGNNIRYIILPDPLALDTLLIDDEPRKKARAARAGASRG
hydra 52 ......................PVQLETLSIRGNNIRYFILPESLPLDTLLVDDEPKSKARGDKGLRGRG
sp_yeast 52 ......................PVPVETLSIRGNNIRYYILPDSLPLDTLLIDDSTKPKQKKKEVVRGRG
sc_yeast 57 SNGIAMASLYLTGGQQPTASDNIASLQYINIRGNTIRQIILPDSLNLDSLLVDQKQLNSLRRSGQIANDP
human_Lsm10 66 ......................QVKLDDLFVTGRNVRYVHIPDDVNITSTIEQQLQIIHRVRN..FG...
human
human 100 RGRGRGR.GR.GRGRGRGGPRR...........
chicken 93 RGRGRGR.GR.GRGRGRGGPRR...........
frog 100 RGRGRGR.GR.GRGRGRGGPRR...........
zebrafish 100 RGRGRGR.GR.GRGRGRGGPRR...........
fruit_fly 100 RGRGRGARGR.GGPRGRGRGRAS.......GRR
worm 100 RGRGGMRGGRGGRGRGRGGPRGGG.....PRR.
hydra 100 RGGP..R.GR.GRGRGRGGPRGGGGGGGGGPGRR
sp_yeast 100 RGRGRGTRGR.GRGASRGF..............
sc_yeast 127 SKKR..RRDF.GAP.ANKRPRRGL.........
human_Lsm10 109 .GKGQGRWEFP........PKNCK.........


C

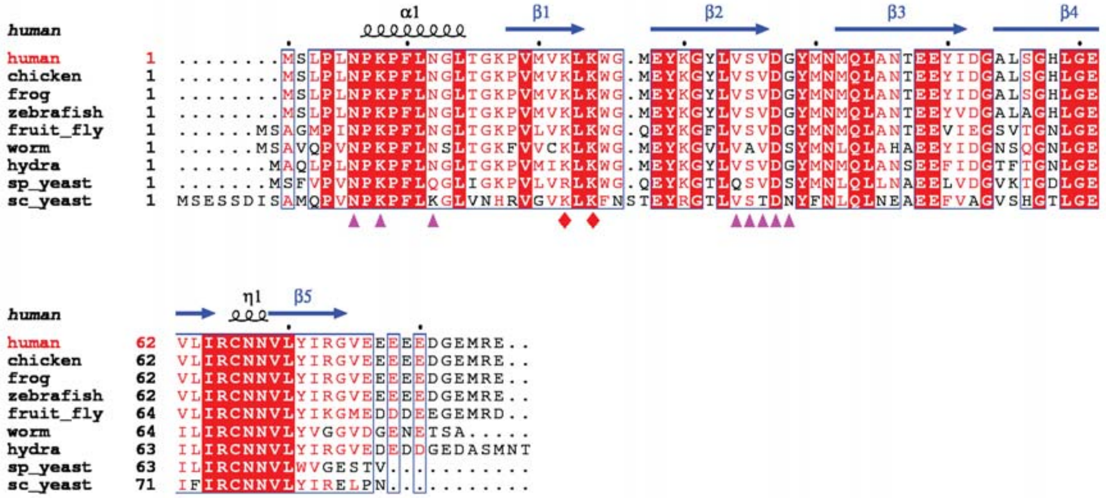

human
α1
β1
β2
β3
β4
human 1 ........MSLPLNPKPFLNGLTGKPVMVKLKWG.MEYKGYLVSVDGYMNMQLANTEEYIDGALSGHLGE
chicken 1 ........MSLPLNPKPFLNGLTGKPVMVKLKWG.MEYKGYLVSVDGYMNMQLANTEEYIDGALSGHLGE
frog 1 ........MSLPLNPKPFLNGLTGKPVMVKLKWG.MEYKGYLVSVDGYMNMQLANTEEYIDGALSGHLGE
zebrafish 1 ........MSLPLNPKPFLNGLTGKPVMVKLKWG.MEYKGYLVSVDGYMNMQLANTEEYVDGALAGHLGE
fruit_fly 1 ......MSAGMPINPKPFLNGLTGKPVLVKLKWG.QEYKGFLVSVDGYMNMQLANTEEVIEGSVTGNLGE
worm 1 ......MSAVQPVNPKPFLNSLTGKFVVCKLKWG.MEYKGVLVAVDSYMNLQLAHAEEYIDGNSQGNLGE
hydra 1 .......MAQLPINPKPFLNGLTGKPVMIKLKWG.MEYKGYLVSVDGYMNMQLANSEEFIDGTFTGNLGE
sp_yeast 1 .......MSFVPVNPKPFLQGLIGKPVLVRLKWG.QEYKGTLQSVDSYMNLQLLNAEELVDGVKTGDLGE
sc_yeast 1 MSESSDISAMQPVNPKPFLKGLVNHRVGVKLKFNSTEYRGTLVSTDNYFNLQLNEAEEFVAGVSHGTLGE
human
η1
β5
human 62 VLIRCNNVLYIRGVEEEEEDGEMRE..
chicken 62 VLIRCNNVLYIRGVEEEEEDGEMRE..
frog 62 VLIRCNNVLYIRGVEEEEEDGEMRE..
zebrafish 62 VLIRCNNVLYIRGVEEEEEDGEMRE..
fruit_fly 64 VLIRCNNVLYIKGMEDDDEEGEMRD..
worm 64 ILIRCNNVLYVGGVDGENETSA.....
hydra 63 ILIRCNNVLYIRGVEDEDDGEDASMNT
sp_yeast 63 ILIRCNNVLWVGESTV...........
sc_yeast 71 IFIRCNNVLYIRELPN...........

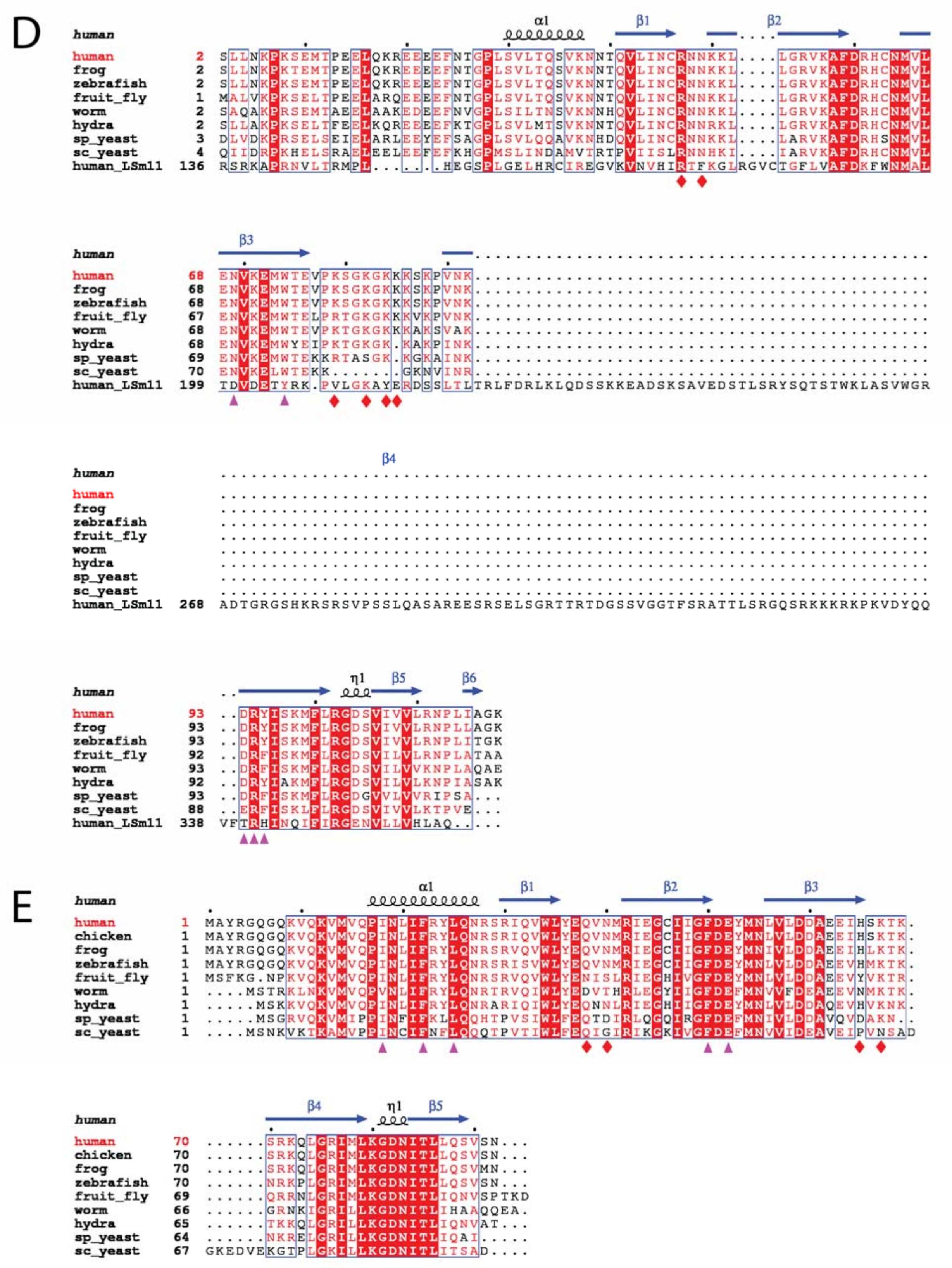
D
E
human
frog
zebrafish
fruit_fly
worm
hydra
sp_yeast
sc_yeast
human_LSm11
chicken

Figure S1.

**Figure S1. Multiple sequence alignments of the orthologues of Gemin2 and SmD1, D2, F and E from diverse organisms.**

Multiple sequence alignment was performed by Clustal Omega [1], and the figures were prepared by ESPript [2]. Absolute identical residues (100%) are in white font with red background. Highly conserved residues (over 70%) are in red font and boxed in blue. The secondary structure of each human protein is shown on the top. Triangles indicate residues interacting between Gemin2 and Sm proteins. Red diamonds indicate residues interacting with 3'-SL RNA.

(A) Alignment of full-length Gemin2 orthologues. The detailed information of species, colors and symbols are the same as in Figure 3.

(B) SmD1 orthologues and human Lsm10. Human (NP_008869); chicken (NP_001264927); frog (NP_001085322); zebrafish (NP_775359); fruit_fly (NP_524774); worm (NP_495306); hydra (XP_002163302); sp_yeast (594613); sc_yeast (NP_011588). human_Lsm10 (NP_116270)

(C) SmF orthologues. Human (NP_003086); chicken (NP_001264291); frog (NP_001080901); zebrafish (NP_001003881); fruit_fly (NP_523708); worm (NP_498708); hydra (XP_002166685); sp_yeast (NP_596101); sc_yeast (NP_015508).

(D) SmD2 orthologues and human Lsm11. Human (NP_004588); frog (NP_001167498); zebrafish (NP_001017582); fruit_fly (NP_649645); worm (NP_506004); hydra (XP_002158231); sp_yeast (NP_594506); sc_yeast (NP_013377). human_Lsm11 (NP_775762)

(E) SmE orthologues. Human (NP_003085); chicken (NP_990581); frog (NP_001085570); zebrafish (NP_957298); fruit_fly (NP_609162); worm (NP_499620); hydra (XP_002154306); sp_yeast (NP_595724); sc_yeast (NP_014802).

A

| RMSD (Å) | SmD1 | SmD2 | SmF | SmE | SmG |
|---|---|---|---|---|---|
| **SmD1** | **0.27** | 0.29 | 0.33 | 0.36 | 0.32 |
| **SmD2** | 0.23 | **0.12** | 0.16 | 0.40 | 0.29 |
| **SmF** | 0.31 | 0.12 | **0.06** | 0.25 | 0.16 |
| **SmE** | 0.29 | 0.23 | 0.20 | **0.14** | 0.17 |
| **SmG** | 0.60 | 0.57 | 0.48 | 0.45 | **0.45** |

B

| RMSD (Å) | SmD1 | SmD2 | SmF | SmE | SmG |
|---|---|---|---|---|---|
| **SmD1** | **0.37** | 0.72 | 0.54 | 0.63 | 0.83 |
| **SmD2** | 0.60 | **0.32** | 0.67 | 0.57 | 0.76 |
| **SmF** | 1.41 | 0.80 | **0.35** | 0.47 | 0.70 |
| **SmE** | 1.27 | 0.83 | 0.48 | **0.41** | 0.48 |
| **SmG** | 1.31 | 1.01 | 0.83 | 0.61 | **0.40** |

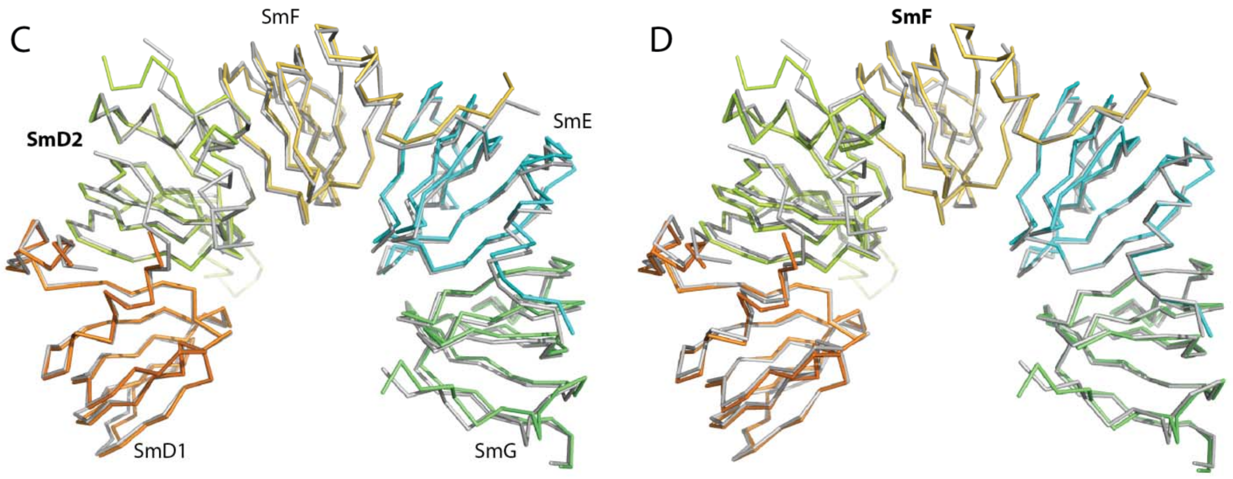


**Figure S2. Sm hetero-pentamer position variations in U1 and U4 snRNP cores.** (A) The RMSDs of Sm proteins from two complexes of U1 snRNP in ASU (PDB code 4PJO) [3] when each pair of Sm proteins is superposed using the 48-residue Sm fold system. (B) The RMSDs of Sm proteins when one U1 snRNP core versus one U4 snRNP core (PDB code 4WZJ) [4]. Superposition of each equivalent Sm protein pair was based on the 48-residue Sm fold system. Superposition of SmD2 (C) and SmF (D) in U1 and U4 snRNP cores are shown in ribbon representation. Sm proteins in U1 snRNP are colored in light gray while Sm proteins D1, D2, F, E and G in U4 snRNP are colored in orange, lemon, yellow, cyan and green respectively.

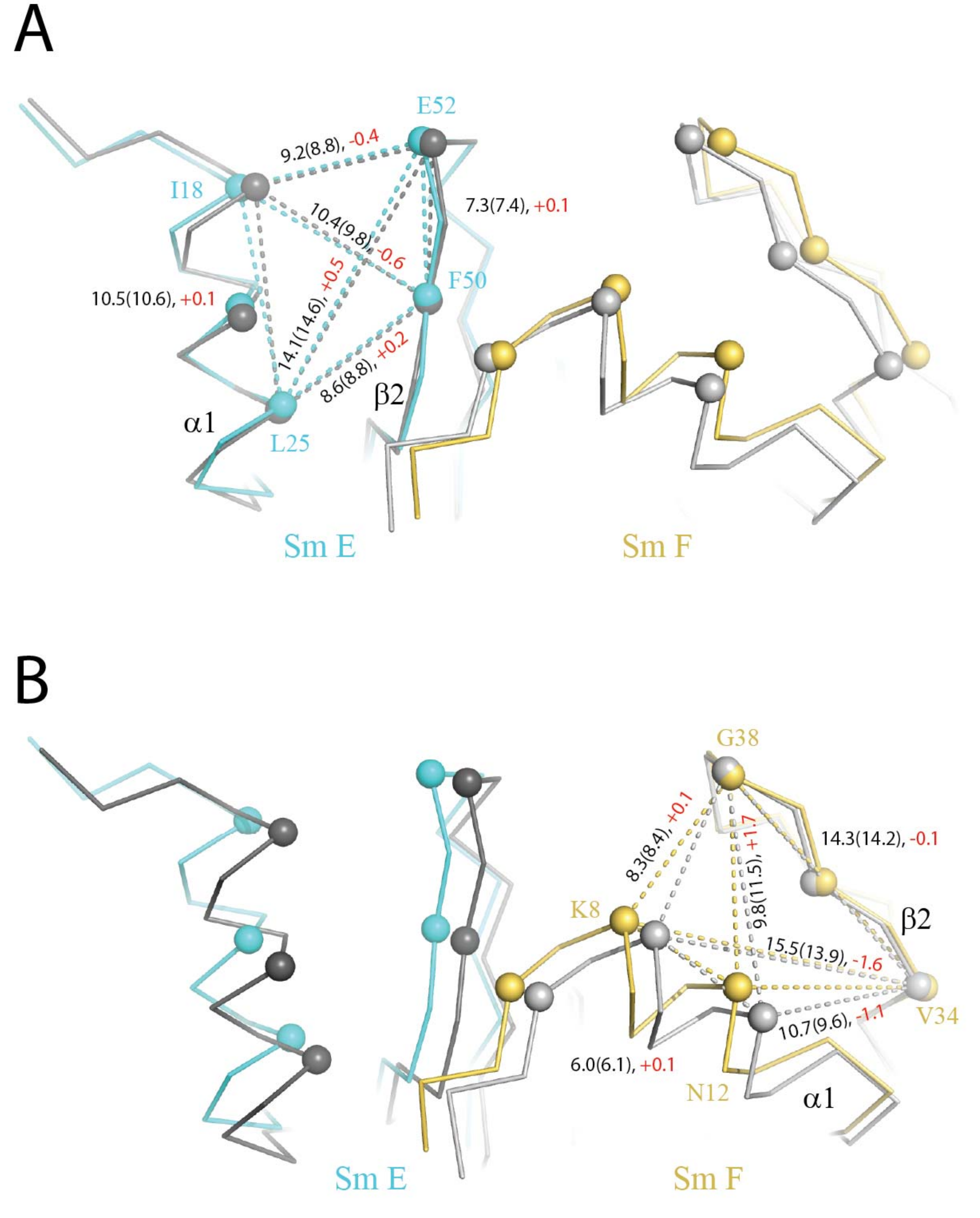


**Figure S3. Conformational changes of Sm F/E from the 7S to the Sm subcore/core.** Cα-Cα distances between the indicated positions on Sm proteins in the 7S and Sm core (U1 snRNP, PDB code 4PJO) [3] are indicated by dashed lines and the distance values (Å) are shown in black with those in the Sm core in brackets. Distance changes are shown in red. SmEs (A) and SmFs (B) in the 7S and Sm core are superimposed based on the 48 positions defined in Figure 6A. SmF/E in U1 snRNP is colored in light gray while SmE and SmF in the 7S complex are colored in cyan and yellow respectively.

**Table S1. Data Refinement Statistics**

| **PDB code** | **3S6N (2011)** | **5XJL (this study)** |
|---|---|---|
| Resolution (Å) | 39.2-2.5 | 39.2-2.5 |
| R factor (%) | 25.3 | 22.1 |
| $R_{free}$ factor (%) | 33.2 | 29.6 |
| Number of reflections for R | 18816 | 18815 |
| Number of reflections for $R_{free}$ | 1015 | 1015 |
| Number of protein atoms | 4553 | 4818 |
| Number of water molecules | 33 | 31 |
| Rmsd bond length (Å) | 0.014 | 0.013 |
| Rmsd bond angles (°) | 1.635 | 1.579 |
| Ramachandran plot (%): | | |
| Favored, additional allowed, disallowed | 92.3, 6.4, 1.3 | 95.8, 2.6, 1.6 |
